# A high-throughput platform for biophysical antibody developability assessment to enable AI/ML model training

**DOI:** 10.1101/2025.05.01.651684

**Authors:** Ammar Arsiwala, Rebecca Bhatt, Lood van Niekerk, Porfirio Quintero-Cadena, Xiang Ao, Adam Rosenbaum, Aanal Bhatt, Alexander Smith, Yaoyu Yang, KC Anderson, Lucia Grippo, Xing Cao, Rich Cohen, Jay Patel, Joshua Moller, Olga Allen, Ali Faraj, Anisha Nandy, Jason Hocking, Ayla Ergun, Berk Tural, Sara Salvador, Joe Jacobowitz, Kristin Schaven, Mark Sherman, Sanjiv Shah, Peter M. Tessier, David W. Borhani

## Abstract

Antibodies must bind their targets with high affinity and specificity to achieve useful therapeutic activity. They must also possess suitable developability properties (e.g., thermostability, solubility, viscosity, polyreactivity) to ensure favorable manufacturing, formulation, and *in vivo* performance. Both binding and developability properties are inherent to a given antibody amino acid sequence. Identification or selection of antibodies possessing suitable binding characteristics is now routine, and *de novo* computational design models, trained on extensive complementarity-determining region sequence and structural data, are rapidly improving.

Developability properties, however, remain difficult to predict largely due to insufficient training data, with empirical testing being heavily used to avoid challenges in late-stage antibody development. To fill this gap, we built a high-throughput antibody developability assay platform designed to generate the large datasets needed to train improved machine learning (ML) models. We optimized and automated known developability assays [Jain et al., 2017], and developed a robust integrated data analytics pipeline. Here we report data on 246 antibodies—representing 106 approved, 135 clinical-stage, and 5 preregistration/withdrawn molecules—across a panel of 10 developability assays, in a “tidy data” format suitable for AI/ML modeling. We used these data to develop an XGBoost [Chen et al., 2016] ML model that better predicts similarity to approved antibodies compared to conventional use of developability warning thresholds. Additionally, we confirm that preliminary predictive models do improve with more training data. Our high-throughput PROPHET-Ab platform enables data generation at the scale needed to develop improved ML models to predict antibody developability.

**Significance:** Successful antibody drugs exhibit important “developability” properties, beyond tight and specific binding to their target, including high expressibility, high stability and solubility, low aggregation propensity, low viscosity, low polyreactivity, and long *in vivo* half-life. Collectively, developability properties predict favorable manufacturing, storage, administration, and safety, and deficiencies in these properties increase risk for clinical failure. Despite progress in developing machine learning models to predict structure and binding, antibody developability models lag, largely due to a lack of sufficiently large training datasets. We have built a high-throughput platform, PROPHET-Ab, that enables data generation at the scale needed to train improved AI/ML models to predict antibody developability.

## Introduction

Antibodies that successfully traverse clinical development to reach licensure and become commercially marketed drugs encompass two fundamental characteristics. First, successful antibodies possess favorable therapeutic target-binding properties, mediated largely by the six hypervariable complementarity-determining regions (CDRs): they bind with high affinity and fast on-rates, usually to the desired epitope, exhibiting high specificity over alternative targets [Chames et al., 2009, Jain et al., 2017, Lu et al., 2020]. Second, they possess several “developability” properties that enable economical production, stable formulation and convenient dosing, and safe clinical use. Developability comprises *in vitro* properties such as high-titer expression in heterologous hosts (e.g., Chinese Hamster ovary [CHO] cells), high solubility and thermostability, and low self-association and viscosity, as well as *in vivo* properties such as low immunogenicity, low non-specific “sticky” binding, and long half-life [Jain et al., 2017, Ahmed et al., 2021, Jain et al., 2023, Jain et al., 2024]. A third characteristic, not universal but nonetheless important to the efficacy of many therapeutic antibodies, is effector function (e.g., complement fixation, antibody-dependent cellular cytotoxicity, and antibody-dependent cellular phagocytosis), mediated largely by the crystallizable fragment (Fc) region and determined in part by the immunoglobulin (Ig) subclass (i.e., IgG1–IgG4) and glycosylation pattern [Vidarsson et al., 2014, Kaneko et al., 2006, Wang et al., 2017, Shields et al., 2006].

Most of these properties are inherent to an antibody given its amino acid sequence. Accordingly, much antibody drug discovery effort is expended in identifying sequences—heavy and light chain pairing, subclass, and especially CDRs—that together exhibit desirable binding, functional, and developability characteristics. Fifty years of advances now enable the routine and rapid generation of antibodies with suitable binding characteristics [Slavny et al., 2024, Pedrioli et al., 2021, Alfaleh et al., 2020] and, increasingly, with the desired functional characteristics. Immunogenicity has also long been recognized as a key optimization parameter [Carter et al., 2024, Sun et al., 2024]. More recently, antibody developability has become widely appreciated as a set of properties requiring optimization in its own right, important to the clinical and commercial success of antibody drugs. Indeed, there is a need for new methods that can identify holistic developability challenges of a given antibody that transcend the mere identification of problematic sequence motifs.

Moreover, there are signs that optimization of binding may actually degrade developability. Shehata et al., 2019 reported that greater antibody specificity, achieved through affinity maturation, correlates *inversely* with three developability properties: polyreactivity, hydrophobicity, and thermal stability. Several other reports suggest that increased hydrophobicity, which is generally required for suitable antigen binding, increases antibody polyreactivity, especially in the case of HCDR2 [Chen et al., 2024, Lecerf et al., 2023]. This latter finding is intriguing in light of the recent observation that immunodominant public antibody responses are driven by germline-encoded binding motifs in CDRs1 and CDRs2 [Shrock et al., 2023], i.e., certain germline sequences may bias an otherwise favorable antibody manifold toward potent binding (after affinity maturation), but inherently poor developability.

Recent advances in machine learning (ML), in particular protein structure prediction and generative protein design [Senior et al., 2020, Jumper et al., 2021, Baek et al., 2021], are positively impacting the *de novo* design of antibody binding characteristics [Ruffolo et al., 2024, Biswas 2025]. These impressive advances have been achieved in large part because the underlying ML models were trained—explicitly or implicitly—on large datasets, including an ever-increasing quantity of high-resolution structural data in the Protein Data Bank [Burley et al., 2019], massive evolutionary sequence variation co-correlations (e.g., EVcouplings) [Marks et al., 2011, Morcos et al., 2011], extensive databases of CDR sequences [Dudzic et al., 2024], and large natural immune repertoires such as the Observed Antibody Space dataset [Olsen et al., 2022].

ML-based design of antibody developability characteristics, in contrast, is in its infancy [Desautels et al., 2024, Biswas 2025]. The selection of antibodies possessing suitable developability characteristics relies on the application of experimentally based empirical cutoffs that outperform simple, biophysics-based *in silico* predictors [Jain et al., 2023]. Despite their prevalent use, however, empirical developability property cutoffs may not effectively categorize antibodies with respect to favorable or unfavorable preclinical and clinical performance.

Because, like binding, developability is inherent to an antibody sequence, and is reflected by its biophysical properties, this *should* be a tractable ML problem. What is lacking, however, is the massive corpus of data needed to train such models. To add to this complexity, standardization by format (e.g., VHH, IgG, VHH-Fc) and assay also remains a challenge despite recent efforts through the use of competitions to develop benchmarks in the field such as AIntibody [Erasmus et al., 2024] and Ginkgo AbDev Competition [Ginkgo Datapoints, 2025].

To fill this data gap, we created a high-throughput (HT), automated antibody developability assessment platform. Our experimental data generation and analytics pipeline, based on known methods [Jain et al., 2017, Makowski et al., 2021, Kohli et al., 2015, Sule et al., 2011, Liu et al., 2014, Kraft et al., 2020] adapted for HT operation and deployed on an innovative physical and computational infrastructure, enables the acquisition and processing of data across 10+ assays with a demonstrated and validated throughput of 1,000s of antibodies per week. Data-related challenges were recently identified as three of the top four barriers to adoption of artificial intelligence (AI)/ML in biopharmaceutical research and development (R&D) [Jelcic et al., 2025]. In fact, non-standard data formatting, i.e., datasets not suitable for AI/ML training and use, was identified as the greatest unmet need [Jelcic et al., 2025]. Widespread use of our antibody developability platform— *PROPHET-Ab: Platform for Reliable Outcome Prediction in High-throughput Evaluation of Therapeutic Antibodies*—will enable the rapid generation of standardized antibody developability data, including for a growing set of diverse antibody formats, at scales needed for training ML models conditioned on *all* characteristics of successful antibody drugs, and will, we believe, help usher in a new era of improved ML-designed antibodies.

## Results and Discussion

### PROPHET-Ab: A High-Throughput Antibody Developability Platform to Expedite Drug Discovery

We designed Ginkgo’s HT automated antibody developability assessment platform to support the generation of data for several applications, including AI/ML model training and drug discovery campaigns (**Figure 1**).

**Fig. 1.**
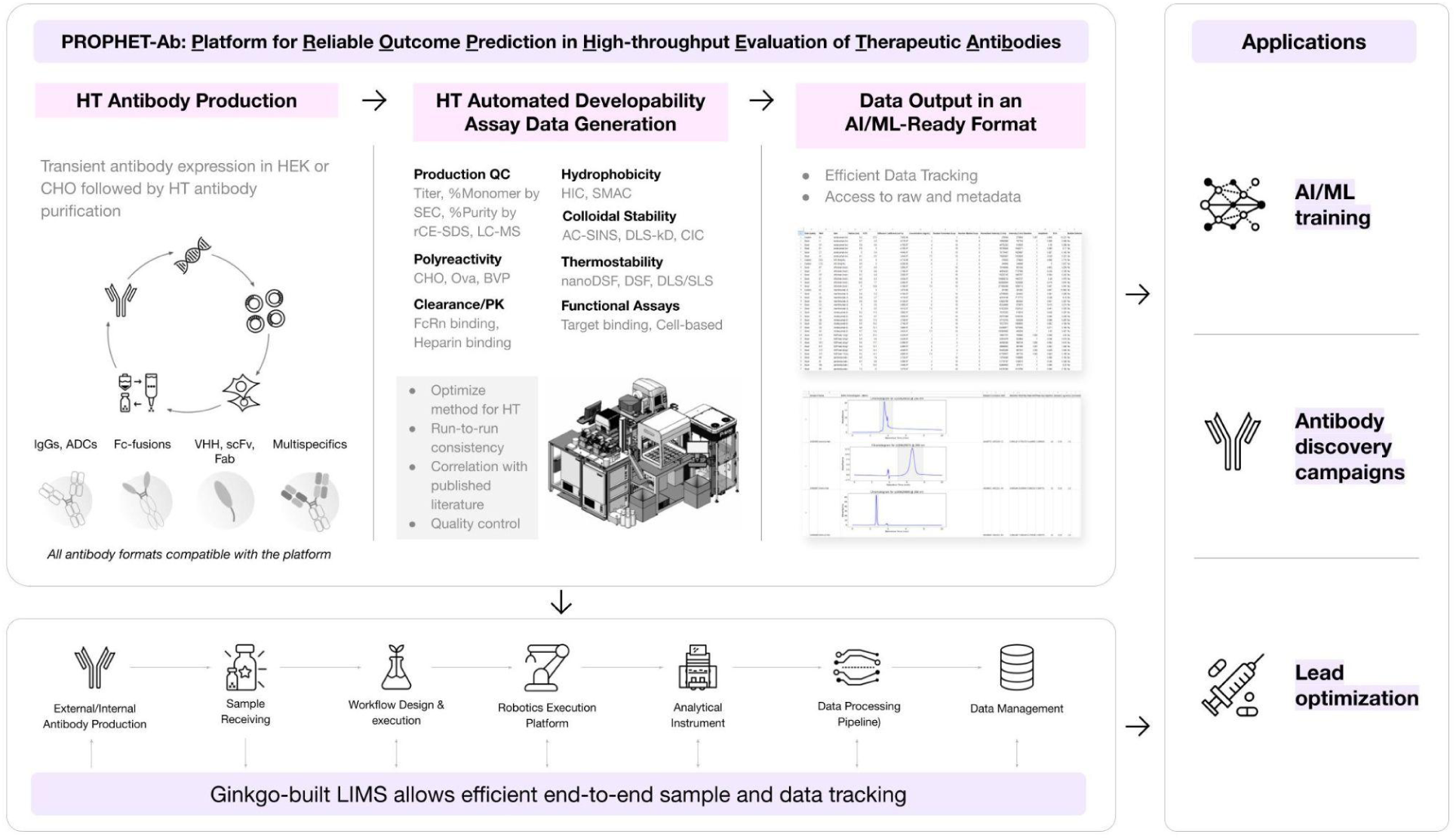
Overview of Ginkgo’s PROPHET-Ab platform for the high-throughput, automated assessment of antibody developability, to support data generation for AI/ML model training and drug discovery campaigns.

The platform is compatible with different antibody formats such as IgGs, antibody-drug conjugates, multispecifics, VHHs, VHH-Fc, minibinders, Fabs, and single-chain variable fragments, and affinity purification tags (e.g., Fc, His, StrepII). Antibodies are transiently expressed in CHO or HEK cells, and purified using the appropriate capture chromatography (e.g., Protein A, Ni-NTA, StrepTactin). Purified antibodies are fed into the characterization engine comprising various standard biophysical assays to assess developability. Data are output in an AI/ML-ready “tidy data” format [Wickham 2014] that facilitates easy access to all raw data and associated metadata. The platform is supported by Ginkgo’s Laboratory Information Management System (LIMS), which enables end-to-end tracking of every sample—from antibody design, through production and purification, automated data acquisition and data processing, to data storage and reporting of results.

### Sequence Characteristics of Antibodies

As a proof of principle, we demonstrate here the utility of the PROPHET-Ab platform to generate developability data for 246 benchmark antibodies. To enable assessment of the impact of variable and constant regions on antibody developability properties, we chose to retain the native IgG subclass originally reported for the antibodies that comprise our test set; all antibodies were grafted on human IgG frameworks. Most were prepared as IgG1s (169); IgG2s (31) and IgG4s (46) round out the set. Kappa light chains (217) predominate, but a few have lambda light chains (29). Antibodies for which the native format is not a standard IgG were prepared by grafting VH and VL domains onto an IgG1 scaffold. For comparison, all the antibodies studied by Jain et al., 2017 were grafted onto an IgG1 scaffold. Sequence, subclass, and immunoglobulin gene information are provided in the “Sequences” data sheet in the supplemental data file.

Our antibody test set spans the entire range from discovery through development to approval and marketing: as of February 2025, 106 are approved for clinical use, while most others are in Phase III (39), II (80) and I (13) clinical trials, and a few are preclinical, withdrawn or have another status (8). Throughout, we compare our results to the Jain et al., 2017 benchmark (henceforth referred to as “Jain dataset”). The results generated using Ginkgo’s HT platform agree well with those using the lower-throughput methods of Jain et al., 2017, Makowski et al., 2021 and Kraft et al., 2020.

### Developability Properties Cluster, in Agreement with Previous Findings

The developability properties (**Figure 2**) assessed in our work comprise expressed titer (mostly in HEK293F cells), % purity (LC+HC [light chain+heavy chain]) measured by reduced, denaturing capillary electrophoresis (rCE-SDS), % monomer (after Protein A purification) by size-exclusion chromatography (SEC), thermostability (T_m1_, T_m2_) by differential scanning fluorimetry (DSF) and nanoDSF, hydrophobicity by hydrophobic interaction chromatography (HIC) and standup monolayer affinity chromatography (SMAC), heparin binding by heparin affinity chromatography (HAC), self-association by affinity-capture self-interaction nanoparticle spectroscopy (AC-SINS) and dynamic light scattering (DLS-kD; DLS-diffusion interaction parameter (kD)), and polyreactivity using the bead-based polyspecificity particle (PSP) method. These assays are summarized in **Table 1**.

**Fig. 2.**
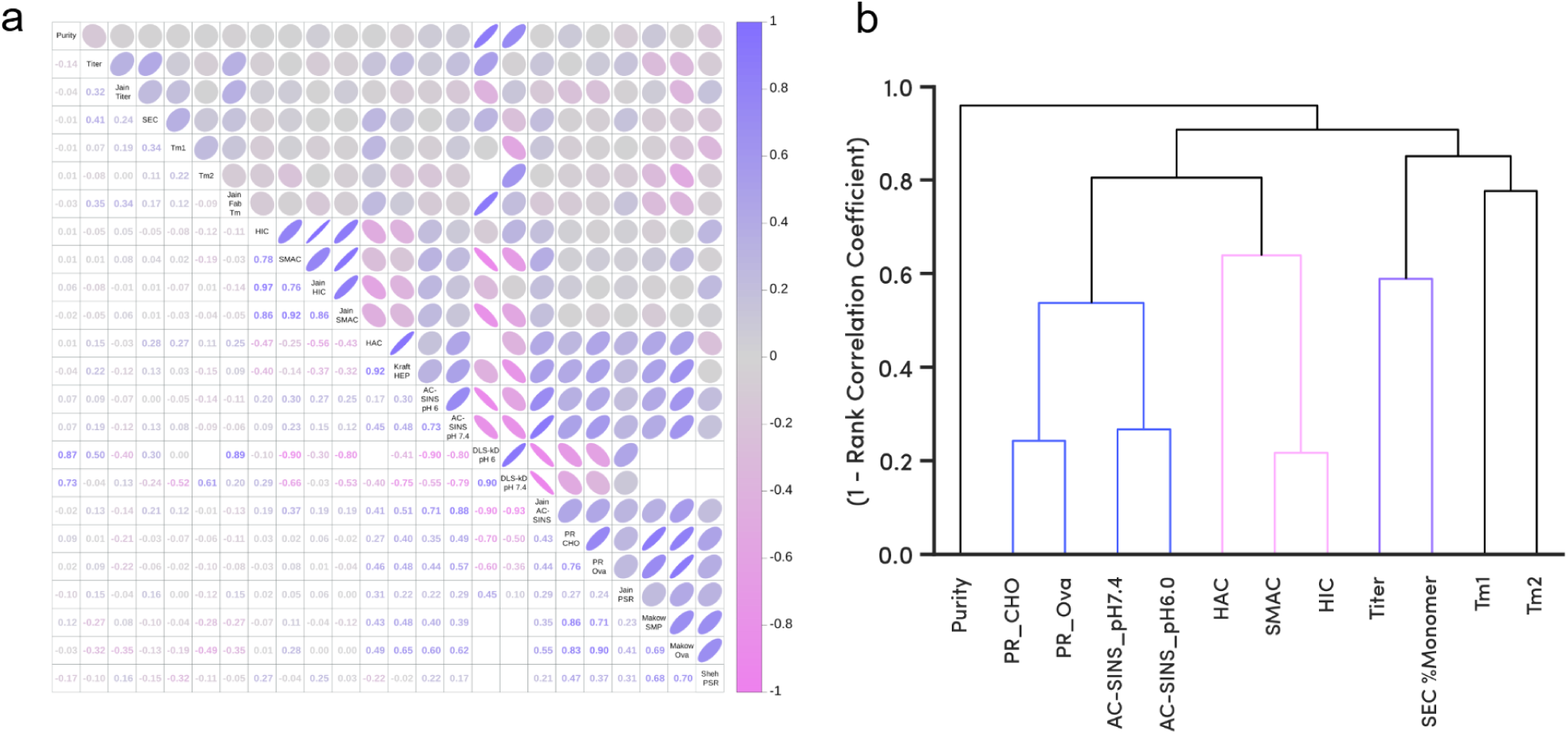
PROPHET-Ab-generated data are consistent with previous findings. (*a*) Cross-correlation of antibody expression, purification, and biophysical properties (both Ginkgo and literature data). Lower triangle: Spearman correlation coefficients; upper triangle: graphical representation. (*b*) Hierarchical clustering of properties (Ginkgo data only; DLS-kD omitted due to few data). Dendrogram scale: property correlation strength, with the most-correlated properties appearing towards the bottom of the dendrogram.

**Table 1.** List of developability assays evaluated in this study and their relevance to manufacturing, formulation, and clinical performance.

| Category | Developability Assay | Readout | Biophysical or Biological Property Measured | Developability Relevance |
| --- | --- | --- | --- | --- |
| Upstream, Expressibility | Titer | mg/L | Expression yield | Composite measure of antibody secretion, folding and correct chain association |
| Purity | CE-SDS | %LC+HC | Fragmentation, Clipping | Impact to potency and increased aggregation |
|  | SE-HPLC | %Monomer | Aggregation | Presence of immunogenic high molecular weight species (HMW) |
| Conformational or Thermal stability | nanoDSF, DSF-SYPRO | Tm1, Tm2 | Temperature induced domain unfolding | Indicative of real-time and accelerated storage stability |
| Hydrophobicity | HI-HPLC | Retention Time (RT) | Surface hydrophobicity | High RT indicative of increased risk of aggregation and non-specific binding |
|  | SMAC | Retention Time (RT) | Surface hydrophobicity, colloidal stability |  |
| Self-Association, Colloidal stability | AC-SINS | $\Delta\lambda_{max}$ | Self-association | Propensity for aggregation, poor colloidal stability and increased viscosity |
|  | DLS-kD | mL/g | Diffusion interaction parameter |  |
| Biological attributes | HAC | Retention Time (RT) | Binding to Heparin | Propensity for non-specific binding to extracellular matrix components. Impact to pharmacokinetics (PK) |
|  | Polyreactivity | PR Score | Binding to CHO lysate or Ovalbumin | Propensity for non-specific off-target binding to non-antigen species. Impact to pharmacokinetics (PK) |

To provide a comprehensive overview of the interrelationships among the various developability properties, and to facilitate comparison of our findings with previously published studies, we generated a correlation matrix and hierarchical clustering dendrogram (**Figure 2**). **Figure 2a** shows the Spearman (rank-order) correlation coefficients between all measured properties, including both Ginkgo-generated data used in this study and previously published data (Jain et al., 2017, Makowski et al., 2021, Shehata et al., 2019, Kraft et al., 2020); **Figure 2b** depicts the hierarchical clustering (Ginkgo-generated data only; note reversed scale). We observe three property clusters with high to moderate correlation coefficients: HIC with SMAC (ρ=0.78); polyreactivity CHO SMP (CHO solubilized membrane protein fraction) with polyreactivity Ova (hen egg white ovalbumin) (0.76); and AC-SINS with DLS-kD [–0.9 (pH 6.0), –0.79 (pH 7.4)].

The polyreactivity and AC-SINS clusters also form a cluster (0.57 to 0.37), as do HIC and SMAC with HAC (–0.47, –0.25). Other properties are less well correlated [e.g., % monomer with titer (0.41)] or not at all, such as purity, in accord with previous findings [Jain et al. 2017].

The cluster including HIC and SMAC reflects that both assays measure the tendency to associate with a hydrophobic column matrix. Both assays correlate well with the Jain dataset, supporting the robustness and consistency of our HT chromatography methods. The good correlation of polyreactivity data generated using CHO SMP versus Ova, and with the Ova data of Makowski et al., 2021, highlights the potential of substituting the simpler Ova for the more complex CHO SMP reagent in early-stage polyreactivity assessment. The weak correlation we observe with polyreactivity data of Jain et al., 2017 and Shehata et al., 2019 is likely due to significant differences in methodology and assay formats. Despite limited data, AC-SINS and DLS-kD, which both assess self-association tendencies, are well correlated (–0.90 to –0.57).

The kD value is a strong predictor of poor solution behavior [Kingsbury et al. 2020, Connolly et al. 2012, Saluja et al., 2010], but this assay has limited throughput. Our findings thus suggest that the AC-SINS assay, with its relatively higher throughput, can be used for rapid, early-stage screening of multiple formulation buffers.

### Purity and Aggregation Depend on IgG Isotype

We mostly used transient transfection in HEK293F cells to generate material for developability testing. Antibodies were produced primarily in two batches; expression titers are shown in **Figure 3a**, along with a comparison to the Jain dataset. As expected for transient expression (Vink et al., 2014), titers averaged ∼200 µg/mL or greater (range 0–800 µg/mL), and appear to be independent of IgG subclass. The titers across these three groups (Jain dataset, Ginkgo Batch 1, Ginkgo Batch 2) cannot be directly compared because of differences in codon optimization, expression vectors, and expression format and conditions.

**Fig. 3.**
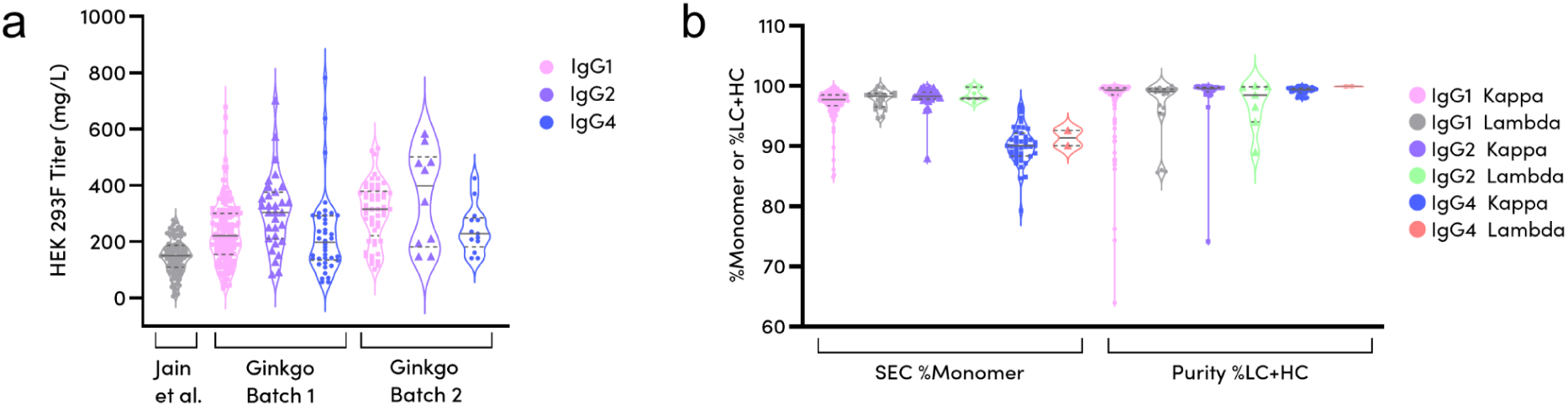
(*a*) Antibody titers (clarified harvest supernatants). Two Ginkgo production batches, colored by IgG subclass, are compared with the results of Jain et al. 2017 (all IgG1). (*b*) Antibody purity, as assessed by aggregation state (%Monomer) and composition (%LC+HC), averaged for batches 1 and 2; colored by IgG subclass and light chain type.

Two measures of antibody purity are shown in **Figure 3b**. The fraction of antibodies present in the expected monomeric state (% monomer), rather than as higher-order aggregates, dissociated chains, or proteolytic fragments, was assessed by SEC. Capillary electrophoresis in the presence of a reducing agent and denaturing detergent, sodium dodecyl sulfate (rCE-SDS), reports on compositional purity—the fractional sum of intact light and heavy chains versus all other protein forms (%LC+HC). Data are shown for 246 IgGs, grouped by subclass (IgG1, IgG2, IgG4) and light chain type (kappa, lambda).

The SEC data indicate that most of the antibodies are monomeric after Protein A purification, with IgG1s and IgG2s generally exhibiting >95% monomer (97.4 ± 2.3%; **Figure 3b**). In contrast, IgG4s exhibit significantly lower % monomer (90.2 ± 3.1%), consistent with the known tendency of this subclass to form both aggregates and fragments (van der Neut Kolfschoten et al., 2007; Kingsbury et al., 2020). rCE-SDS data confirm the high overall purity across all subclasses, indicating robust expression and assembly. A few IgG1/2 kappa and lambda antibodies, however, do exhibit fragmentation, as shown by the long tails toward lower %LC+HC. These observations highlight the importance of considering trade-offs between antibody developability and choice of IgG subclass during candidate selection and preclinical development. IgG1s and IgG2s possess effector function and have higher % monomer, but some also exhibit greater fragmentation, and thus a potentially more complex or lower yielding purification process. Conversely, IgG4s have low effector function, often a desirable characteristic, but their inherent biophysical properties can affect overall homogeneity and stability, requiring tailored optimization strategies during development.

### IgG4s Exhibit Lower Thermal Stabilities than Other Subclasses

Thermostability profiles of the 246 antibodies, determined by nano differential scanning fluorimetry (nanoDSF) that monitors intrinsic tryptophan/tyrosine fluorescence, are presented in **Figure 4a**. The data, grouped by heavy chain (IgG1, IgG2, IgG4) and light chain (kappa, lambda) types, reveal distinct thermal unfolding patterns. IgG4s generally exhibited a lower median T_onset_ and T_m1_, compared to IgG1s and IgG2s, potentially indicating reduced overall stability and increased conformational flexibility [van der Neut Kolfschoten et al., 2007]. T_m1_ values, which typically represent CH2 domain unfolding, were relatively consistent across all IgG subclasses, suggesting similar stability of this half of the Fc region. T_m2_ values, which represent Fab and CH3 domain unfolding, did reveal some variability. Some IgG4 kappa antibodies exhibited notably higher T_m2_ values compared to other subclasses, possibly indicating a more stable Fab domain in this particular subclass and light chain combination. Our data suggest that the IgG subclass, and to a lesser extent the light chain type, can significantly impact antibody thermal stability profiles, with IgG4s generally displaying a lower onset of unfolding and potentially more stable Fab regions in specific cases [Ionescu et al., 2008, Kim et al., 2022].

**Fig. 4.**
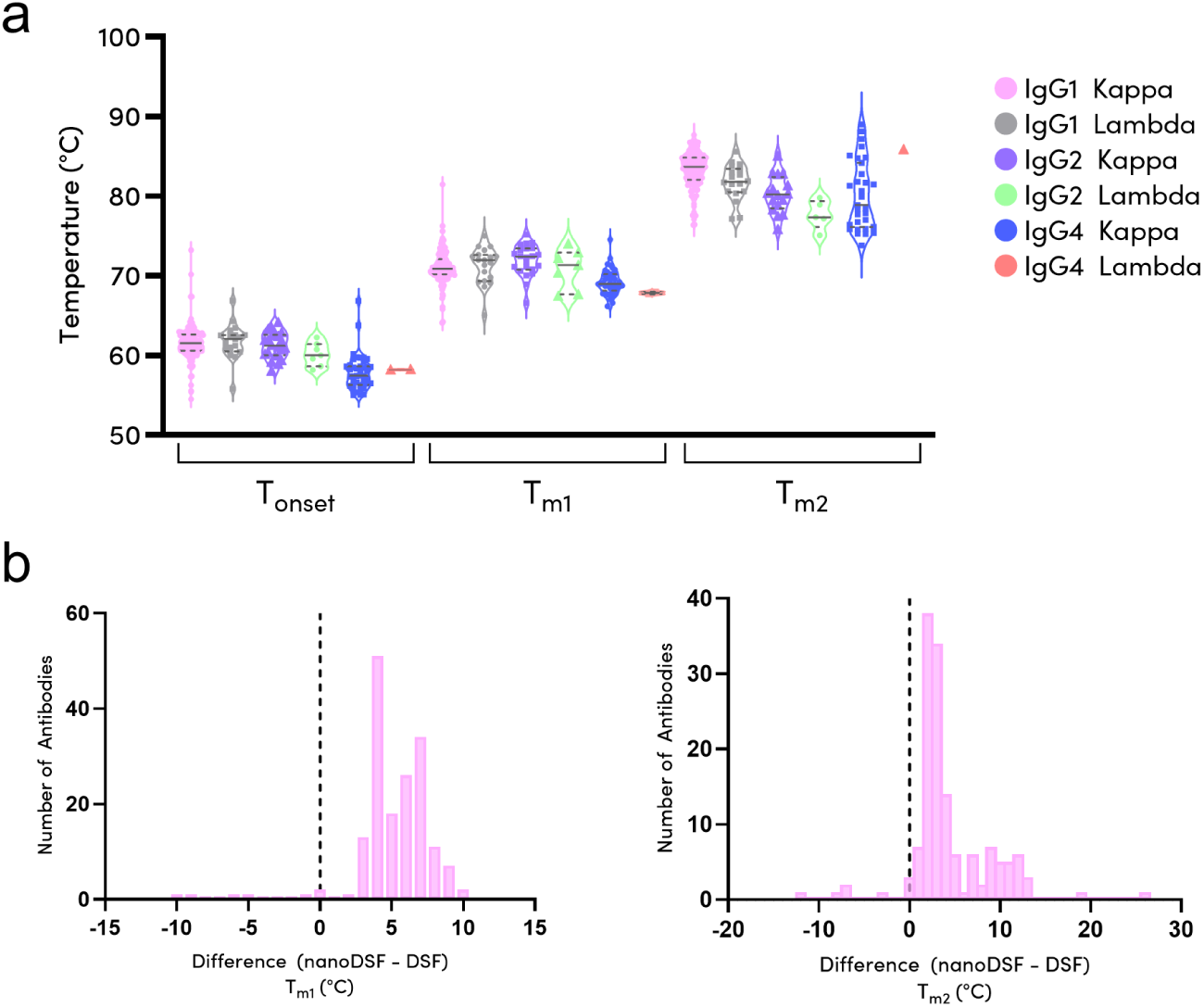
*(a)* IgG subclasses exhibit different thermal stabilities. Melting temperatures determined by nanoDSF (246 IgGs). *(b)* Dye-based DSF systematically underestimates T_m1_ and T_m2_ compared to nanoDSF (174 IgGs).

We also compared thermostability data obtained using nanoDSF with traditional dye-based DSF for a subset of 174 IgGs (data in the ‘nanodsf vs dsf’ sheet in the supplemental data file). The selection of the 174 IgGs for dye-based DSF analysis was based on sample availability; samples with limited yield were not assayed. We observed a systematic deviation, with nanoDSF consistently reporting higher melting temperatures (**Figure 4b**). Our observations align with previous reports demonstrating that dye-based DSF methods can underestimate the true protein stability, as interaction of proteins with hydrophobic dyes appears to facilitate domain unfolding, thereby slightly lowering the observed melting temperature [Magnusson et al., 2018, Gao et al., 2020]. In contrast, nanoDSF, which relies on the intrinsic fluorescence of tryptophan and tyrosine residues, provides a label-free, non-disruptive measure of protein stability [Magnusson et al., 2018], potentially offering a more accurate representation of the inherent thermostability of proteins. It is also important to note that the choice of buffer and pH can significantly influence the melting temperatures and the nature of the unfolding transitions observed [Kim et al., 2022]. The use of intrinsic fluorescence also has limitations, such as the lower sensitivity of the method, compared to dye-based DSF methods, at lower antibody concentrations.

Although determination of Fab thermostability is an important part of early-stage developability assessments, purification of Fabs from thousands of IgGs for independent analysis is impractical. As well, nanoDSF or DSF do not typically resolve the three unfolding transitions (Fab, CH2, CH3) expected of an intact IgG: Fab unfolding frequently overlaps either CH2 or CH3 unfolding. The inability to deconvolute with high confidence the Fab, CH2, and CH3 unfolding transitions when using a full-length IgG as the input sample, rather than an Fab, likely explains the weak correlation between the T_m1_/T_m2_ data from our study (nanoDSF and dye-based DSF) compared to the Fab T_m_ reported in the Jain dataset. Additionally, the buffer background and ramp rates between the two studies were also different, likely weakening the correlation further. These considerations highlight a gap in our current ability to reliably determine Fab T_m_ values using standard DSF data, hindering the full utilization of this HT method for early-stage developability assessment. This challenge, and potential approaches to resolve it through advanced data analysis and deconvolution techniques, will be addressed in future work.

### High-Throughput Chromatography Assays Correlate Strongly with Prior Methods

Assays that rely on chromatographic separation are generally not suited for HT operations and have historically limited the generation of antibody developability data. We have developed HT methods for assessing hydrophobicity (HIC), colloidal stability (SMAC), and heparin binding (HAC) by modifying and optimizing literature methods [Jain et al., 2017, Kohli et al., 2015, Kraft et al., 2020]. Our improved methods offer shorter run times (∼5-10 mins/run) without compromising data resolution or quality. **Figure 5** presents a comparative analysis of the data generated using our HT methods with previously published datasets [Jain et al., 2017, Kraft et al., 2020].

**Fig. 5.**
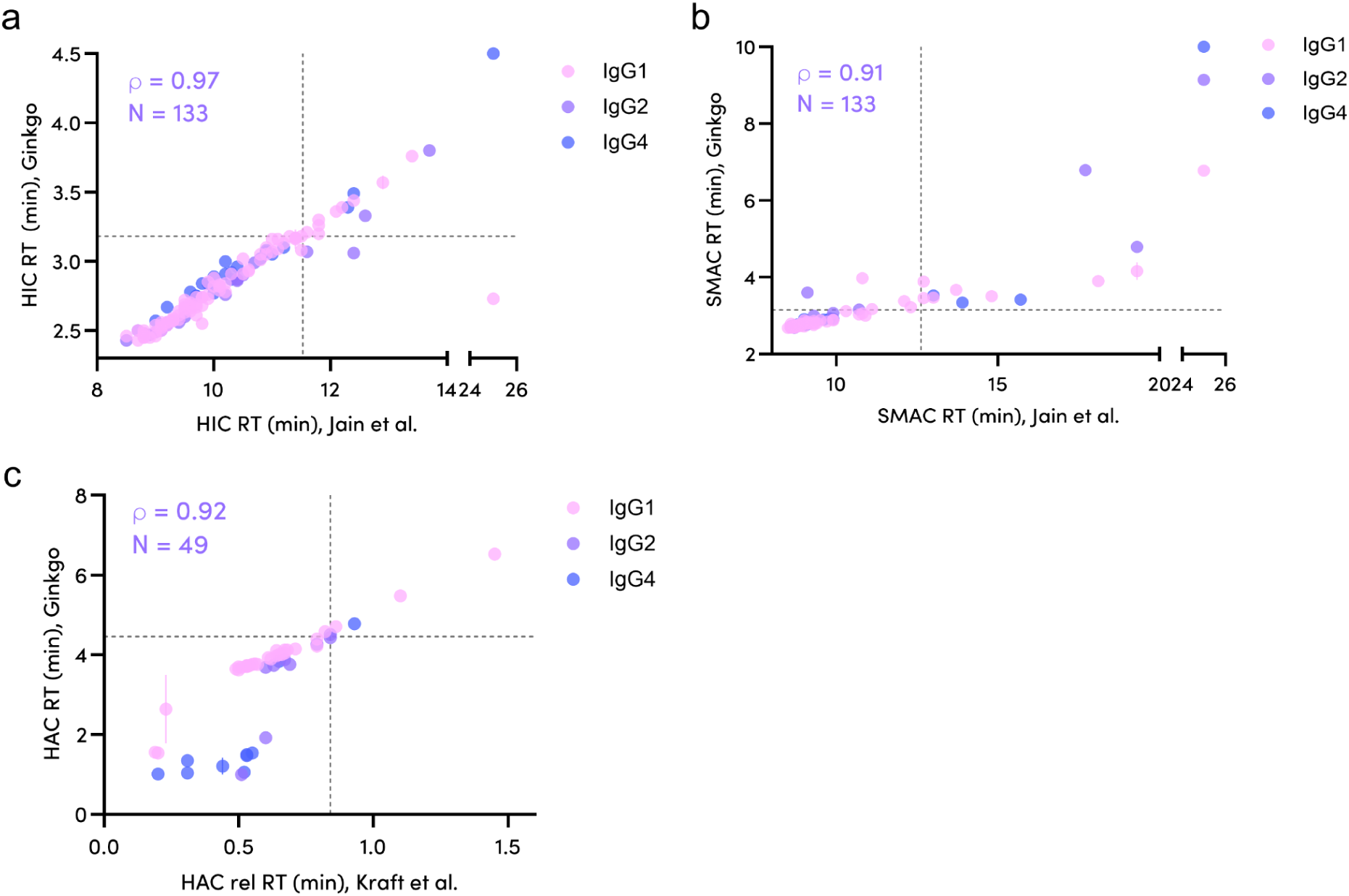
PROPHET-Ab HT chromatography methods generate data comparable to reported lower-throughput methods. *(a)* Hydrophobicity, *(b)* colloidal stability, and *(c)* heparin binding Ginkgo-generated data correlated with data from Jain et al. 2017 (a, b) and Kraft et al. 2020 (c). Warning thresholds are shown in dotted lines (per Jain et al. 2023 or Ginkgo, respectively).

HIC retention times generated using our platform are highly correlated (ρ=0.97, n=133) with those reported in the Jain dataset (**Figure 5a**). Notably, Jain et al. used an IgG1 scaffold for all antibodies. Thus, the observed correlation suggests that interaction with the HIC matrix is largely independent of the IgG subclass, i.e., antibody hydrophobicity is largely a function of the variable regions. This finding is consistent with the understanding that most hydrophobic patches responsible for undesirable hydrophobicity reside within the CDRs [Tang et al., 2021].

Similar to HIC, we observed a strong correlation (ρ=0.91, n=133) between Ginkgo’s SMAC data and those in the Jain dataset (**Figure 5b**). Our improved SMAC method enables a second column volume wash while maintaining a run time >2x faster than the published method (Kohli et al., 2015). This additional wash allows resolution of “sticky” antibodies, potentially improving the discriminatory power of the assay. For instance, our method is able to resolve three highly sticky monoclonal antibodies (mAbs), lirilumab, glembatumumab, and eldelumab, whereas all three eluted together at the end of run (25 min) in the previous method (Kohli et al., 2015). As with HIC, the SMAC correlation appears to be independent of IgG subclass, suggesting that the factors governing SMAC retention are primarily driven by variable region characteristics.

During method development, we also compared our shorter-run-time HIC and SMAC methods with the reported longer-run-time methods (Jain et al., 2017), all run at Ginkgo, for 30 IgGs that were selected as a ladder spanning the range of hydrophobicities based on the Jain dataset.

The strong observed correlations (0.99, 0.96; **Figure S1**) give confidence that our HT methods report faithfully on these key parameters.

Kraft et al., 2020 proposed using heparin as a surrogate for the highly negatively charged glycocalyx components on endothelial cells, among the main contributors to nonspecific antibody clearance. By directly correlating HAC retention time with clearance, they identified HAC as a useful tool to assess differences in nonspecific cell-surface interaction and the likelihood for increased pinocytotic uptake and degradation. Our HAC method has a >3x faster run time than the previous method. For IgG1s, we observe a good correlation between our HAC retention times and those previously reported (**Figure 5c**). The weaker correlations observed for IgG2s and IgG4s are likely attributable to the fact that the previous study grafted all antibodies on an IgG1 framework, and the corresponding differences in charge and glycosylation profiles within the constant regions of these subclasses.

### AC-SINS is a Potential High-Throughput Surrogate for DLS-kD to Assess Self-Association

We used AC-SINS and DLS-kD to assess antibody self-association in two conditions (phosphate-buffered saline (PBS), pH 7.4; histidine/arginine, pH 6.0). AC-SINS (0.025 mg/mL mAb) was performed for all 246 IgGs whereas DLS-kD (1–10 mg/mL mAb) was performed for 10 IgGs. The data (**Figure 6**) highlight the utility of these assays for early developability assessment and formulation screening.

**Fig. 6.**
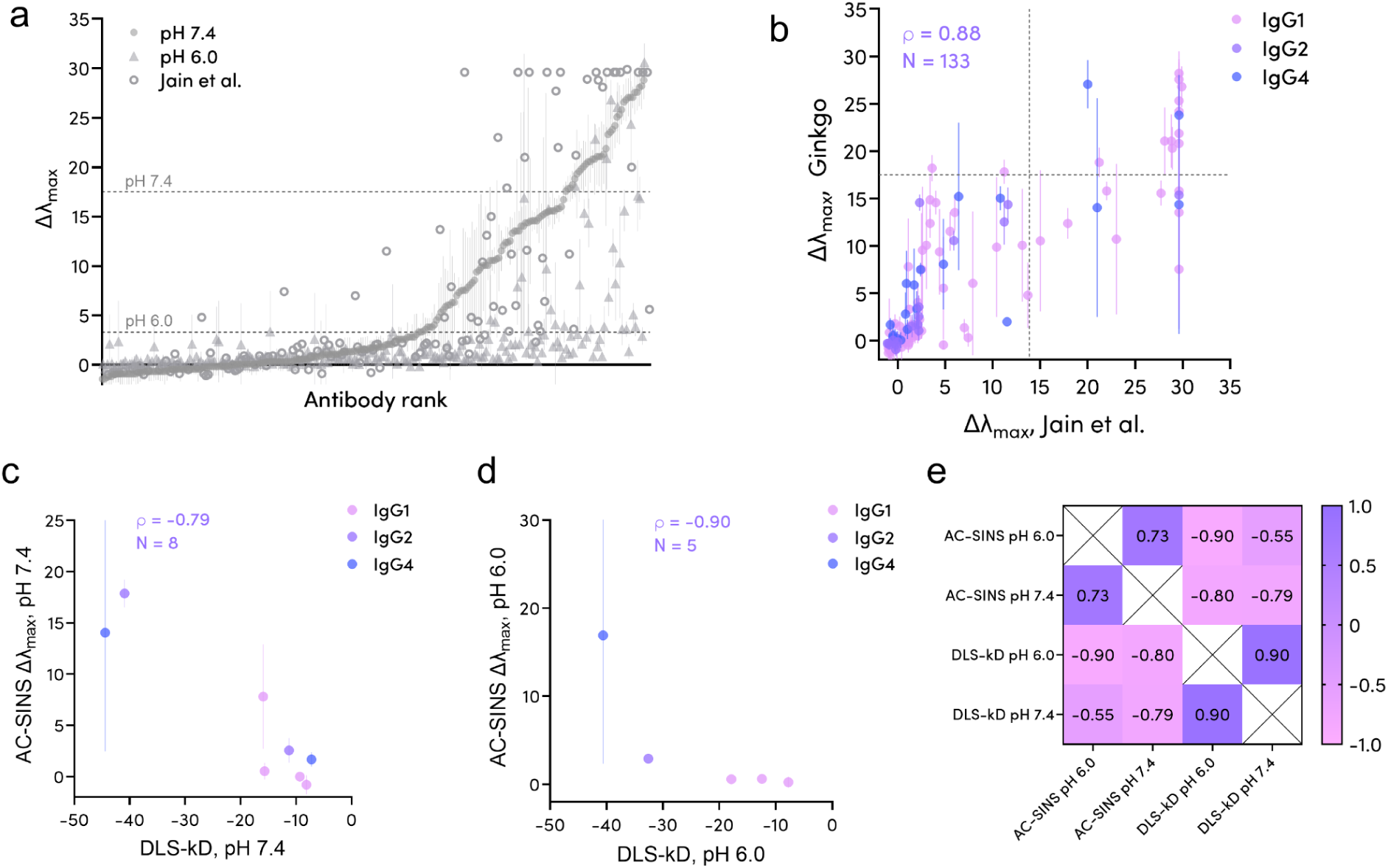
Self-interaction measured by AC-SINS correlates well with DLS-kD. (a) Rank-ordered AC-SINS Δλmax for 246 IgGs in two buffers compared to data reported by Jain et al. 2017. Updated warning thresholds for His/Arg, pH 6.0 (3.3) and PBS, pH 7.4 (17.51) are shown as dotted lines. (b) Correlation of Ginkgo (PBS, pH 7.4) with Jain et al. 2017 AC-SINS Δλmax data. Warning thresholds are shown as dotted lines (Jain et al. 2023, 13.84; Ginkgo, 17.51). (c, d) Correlation of Ginkgo DLS-kD with AC-SINS Δλmax data (PBS pH 7.4; His/Arg, pH 6.0). (e) Spearman rank correlation matrix between AC-SINS (for 246 Abs) and DLS-kD (for 10 Abs), each in two buffers.

We rank-ordered antibodies by AC-SINS Δλmax value in PBS at pH 7.4 (solid circles); higher Δλmax values correspond to increased self-association (**Figure 6a**). Antibodies tended to exhibit a lower Δλmax, i.e., lower self-association, in histidine/arginine buffer at pH 6.0 (solid triangles), presumably due to increased electrostatic repulsion. The presence of arginine and polysorbate-80 likely further contributes to shielding self-interaction. The downward shift in Δλmax when going from pH 7.4 to pH 6.0 (a typical formulation buffer pH) highlights the benefit of using HT methods such as AC-SINS for early formulation screening.

Our AC-SINS data (at pH 7.4) correlate well with those reported in the Jain dataset (**Figure 6b**; ρ=0.88, n=133). The data mostly cluster along the diagonal, indicating a good agreement between the two datasets and validating the reliability of our HT AC-SINS platform.

Using a panel of 29 antibodies (28 IgG1s, 1 IgG4), Connolly et al., 2012 demonstrated that the antibody DLS-kD is a useful predictor of viscosity in both high and low ionic strength buffer systems. Kingsbury et al., 2020 expanded on this study by measuring DLS-kD for 59 mAbs (44 IgG1s, 4 IgG2s, 11 IgG4s) and correlating the measurements with the high concentration opalescence and viscosity behavior of the mAbs. **Figures 6c** and **6d** show correlations between AC-SINS Δλmax and DLS-kD in PBS, pH 7.4 (n=8) and histidine/arginine, pH 6.0 (n=5), respectively. The sample set for this correlation between AC-SINS and DLS-kD is admittedly small. However, the observed rank-order agreement between AC-SINS and DLS-kD is encouraging and an area of expansion for future work. While DLS-kD is a well-established predictor of solution behavior and high-concentration formulation success, its relatively low throughput and requirement for large quantities of purified antibody (1–10 mg/mL), compared to AC-SINS (0.025 mg/mL), limit its application in early-stage developability screening.

The relationships between AC-SINS and DLS-kD data in the two buffer conditions is summarized by a Spearman rank correlation heatmap (**Figure 6e**). A previous study by Mieczkowski et al.. 2021 showed a parabolic relationship between DLS-kD and AC-SINS measurements. Therefore, while our work is not the first to show a correlation between these methods [Zarzar et al. 2023, Connolly et al. 2012, Kingsbury et al. 2020], the strong positive correlations observed between AC-SINS and DLS-kD across both pH 7.4 and pH 6.0 buffer conditions further support the value of AC-SINS, particularly for the rapid screening of multiple formulation buffers in parallel.

### Polyreactivity Assessment Using CHO Solubilized Membrane Protein and Ovalbumin Are Comparable

We assessed IgG polyreactivity against solubilized membrane proteins (SMP) from CHO cells and ovalbumin (Ova), which are complex and simple polyreactivity reagents, respectively. The data shown in **Figure 7** highlight the performance and potential advantages of the bead-based polyreactivity assay originally proposed by Makowski et al. 2021, which we have onboarded on automated workcells for increased throughput and reliability.

**Fig. 7.**
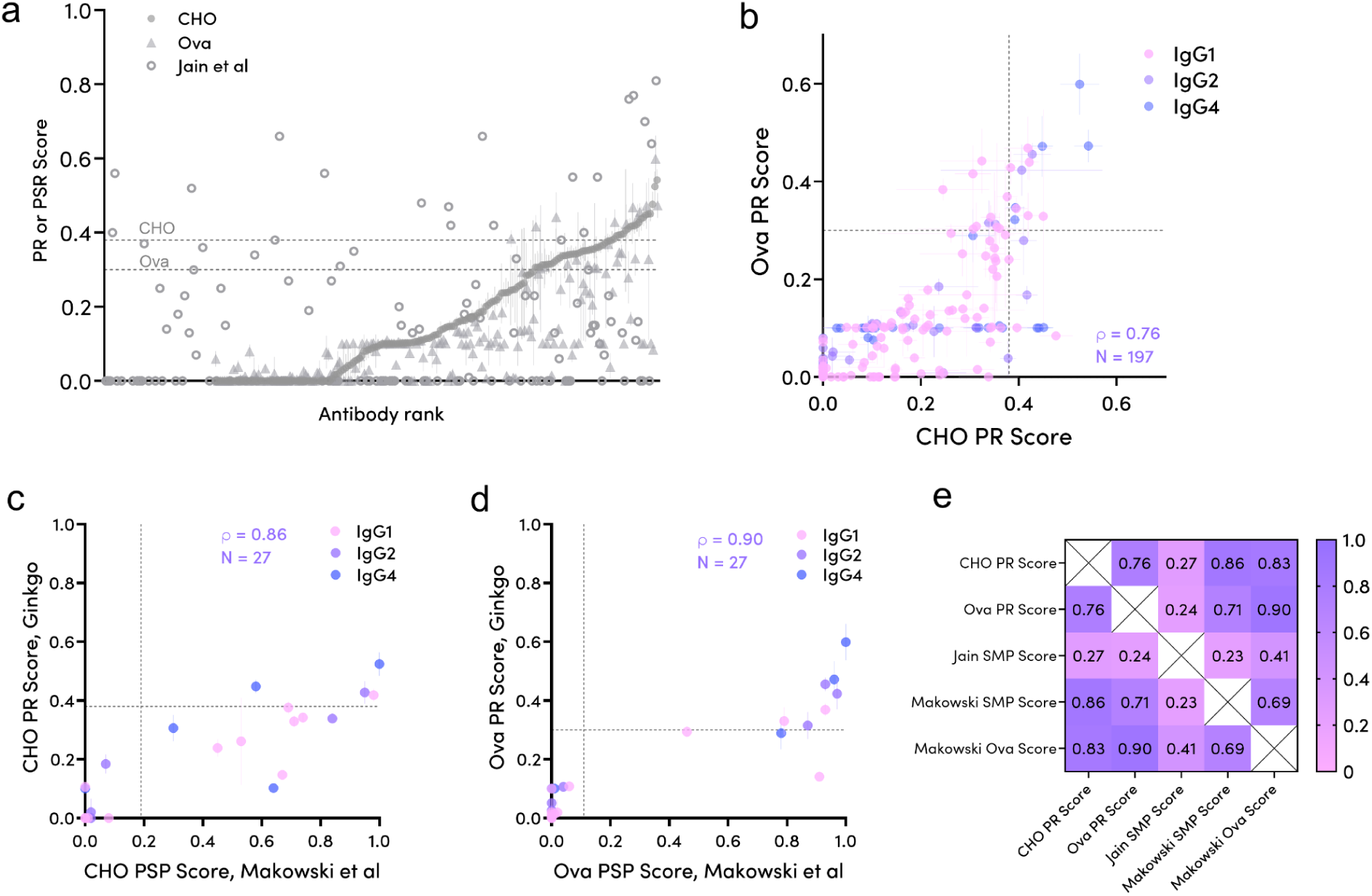
Polyreactivity (PR) scores against CHO SMP and ovalbumin (Ova) are comparable. *(a)* Rank-ordered PR scores, comparing Ginkgo CHO SMP and Ova with data reported by Jain et al. 2017. Ranked by increasing Ginkgo CHO PR score. Updated warning thresholds for CHO PR (0.38) and Ova PR (0.30) scores are shown as dotted lines. *(b)* Correlation of Ginkgo CHO and Ova PR scores, colored by IgG subclass. Warning thresholds are shown. *(c,d)* Correlation of CHO or Ova PR scores with Makowski et al. 2021 PSP scores. Warning thresholds are shown (horizontal: Ginkgo; vertical: Makowski, but calculated in a manner different from Ginkgo). *(e)* Spearman rank correlation matrix.

Figure 7a shows rank-ordered polyreactivity scores against CHO SMP (solid circles), with the corresponding scores against Ova overlaid (solid triangles); scores were normalized to control antibodies (elotuzumab, dalotuzumab, ixekizumab) using the method proposed by Shehata et al.. 2019. Our data generally show good agreement between CHO SMP and Ova (Figure 7b), but do not align well with the proprietary yeast-based data in the Jain dataset (**Figure S2a**). We believe this lack of alignment is due to two factors. The Jain dataset was generated using yeast-displayed antibodies, and SMP binding thus involved interaction with antibodies bearing non-mammalian, high-mannose, non-sialylated glycans, as well as with adventitious yeast surface proteins. By contrast, we used purified antibodies expressed in CHO/HEK cells attached via Protein A to passivated Dynabeads.

Polyreactivity scores assessed using CHO SMP versus Ova are presented in Figure 7b; dotted lines mark the thresholds for “flagged” (problematic) antibodies. As expected, we see good agreement between CHO SMP and Ova data, with most antibodies falling into the good (lower-left) or bad (upper-right) quadrants. Expanding on the original data presented by Makowski et al. 2021 (n=32), we present a comparison of the two polyreactivity reagents for a much larger number of antibodies (n=197) spanning different IgG subclasses. Our data suggest that Ova, used as a proxy for CHO SMP in an initial screen, yields similar warning for polyreactive antibodies. Consistent with Makowski’s observations, the bead-based method offers several advantages over the polyspecificity reagent method, including a lower required antibody concentration (≤15 μg/mL), especially useful during early-stage development.

Moreover, the bead-based method is automatable (1,000s of samples/day) and amenable to different Fc-based antibody formats such as VHH-Fc and multi-specific antibodies.

Ginkgo CHO and Ova polyreactivity scores correlate well (ρ=0.86, 0.90) with those reported by Makowski et al., 2021 for a subset of 27 common antibodies (Figure 7c**,d**). Of note, Makowski et al. used an IgG1 framework for all antibodies, whereas we retained the original IgG subclass. The good correlations suggest minimal effect of the Fc region on polyreactivity, again indicating that the most polyreactive regions are predominantly located in the CDRs [Chen et al., 2024]. A Spearman correlation heatmap (Figure 7e) emphasizes the good agreement between our data, collected using either polyreactivity reagent, and Makowski’s bead-based data, but a lack of correlation with Jain’s yeast-based method.

### Developability Assay Sensitivity to Expression Host

To determine the impact of expression host on antibody developability profiles, we expressed 20 IgGs in ExpiCHO cells and subjected them to the same developability assessment panel as performed with HEK293-expressed antibodies. The chosen set of diverse therapeutic antibodies exhibit a spread in heavy and light chain subtypes as well as reported developability properties from the Jain dataset [Jain et al., 2017]. These antibodies were adalimumab, belimumab, bimagrumab, daclizumab, denosumab, evolocumab, farletuzumab, foralumab, glembatumumab, infliximab, mavrilimumab, natalizumab, nimotuzumab, ofatumumab, ozanezumab, pembrolizumab, tovetumab, tralokinumab, vedolizumab, and veltuzumab.

Developability assessment data of HEK and CHO-expressed antibodies are shown in Figure 8. Strong correlations are observed for HIC (r=0.99), AC-SINS pH 6.0 (0.96), PR Score CHO (0.89), T_m1_ (0.77), % monomer (0.75), T_onset_ (0.71), and AC-SINS pH 7.4 (0.68), suggesting that these assays are relatively robust and less sensitive to the host expression system. In contrast, weaker correlations were observed for titer (∼0.51). The poor titer correlations may be attributable to known differences in HEK293 and CHO secretory pathways [Malm et al., 2022].

**Fig. 8.**
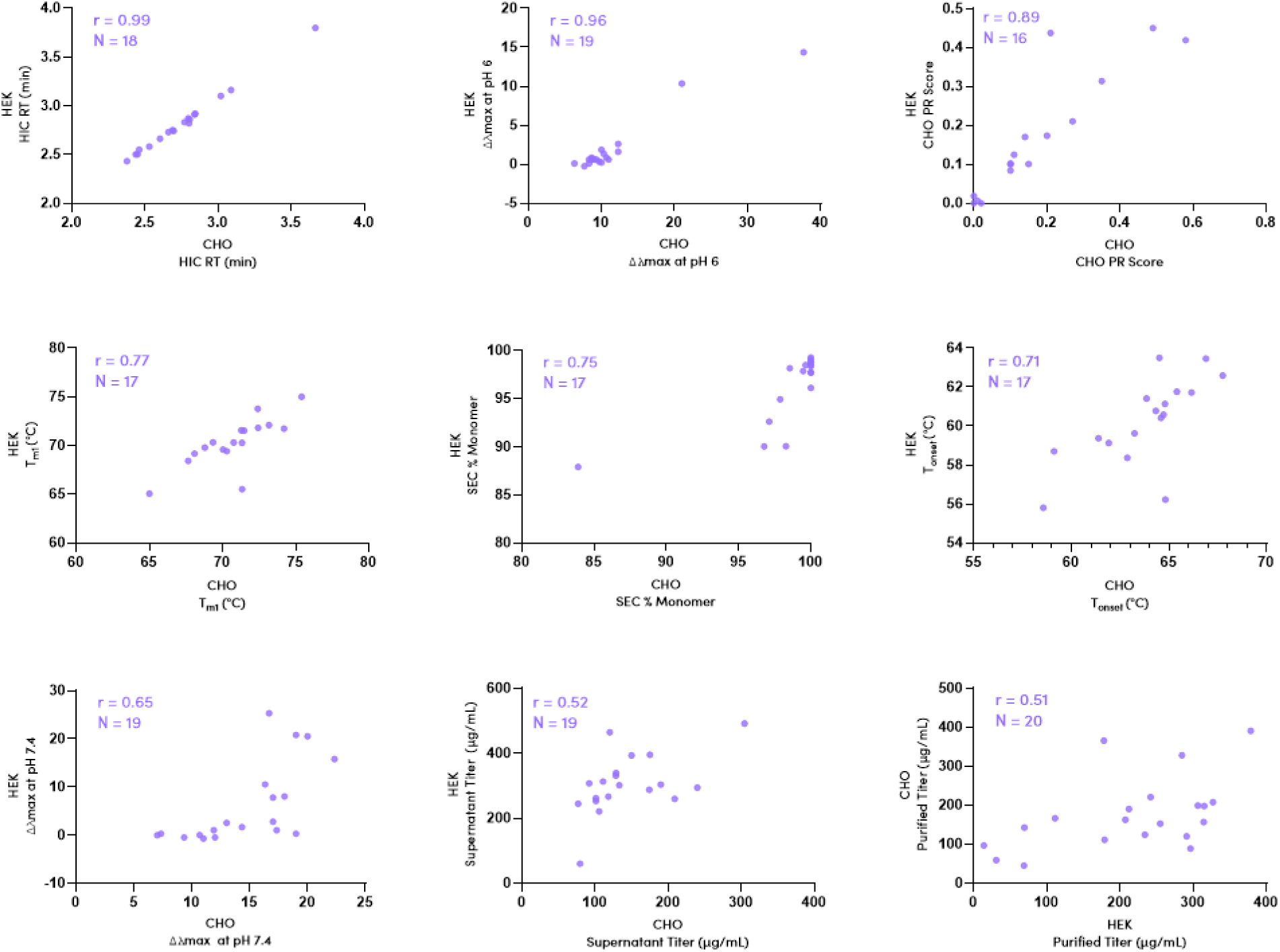
Comparison of HEK-and CHO-produced antibodies (up to 20 IgGs) across nine developability readouts: Hydrophobicity (HIC RT), Self-association (Δλmax pH 6.0), Polyreactivity (CHO PR), Thermostability (T_m1_), Aggregation (SEC %Monomer), Thermostability (T_onset_), Self-association (Δλmax pH 7.4), Supernatant Titer (µg/mL), Normalized Titer (µg/mL).

Our comparative data highlight that key developability properties involving self-interaction and stability are largely independent of the expression host. Thus, use of a convenient expression host (HEK293) for large-scale, early developability assessments of antibody libraries appears to be compatible with later transition to CHO for clinical trial-and commercial-scale production.

### Updated Developability Warning Thresholds Afforded by a Larger Dataset

To refine developability risk assessments, we updated the developability warning thresholds, previously established by Jain et al., 2017 and Jain et al., 2023, using our data on 106 approved antibodies. An antibody is given a warning flag for each of its properties that exceeds a given assay’s threshold. Our updated thresholds (**Table 2**)—the 90th percentile of each assay’s data histogram (Figure 9)—generally lie within the margin of error of the earlier thresholds but exhibit tighter standard deviations, reflecting the increased statistical power afforded by our larger dataset. These updated thresholds provide a refined benchmark for identifying antibody candidates at risk of poor developability.

**Fig. 9.**
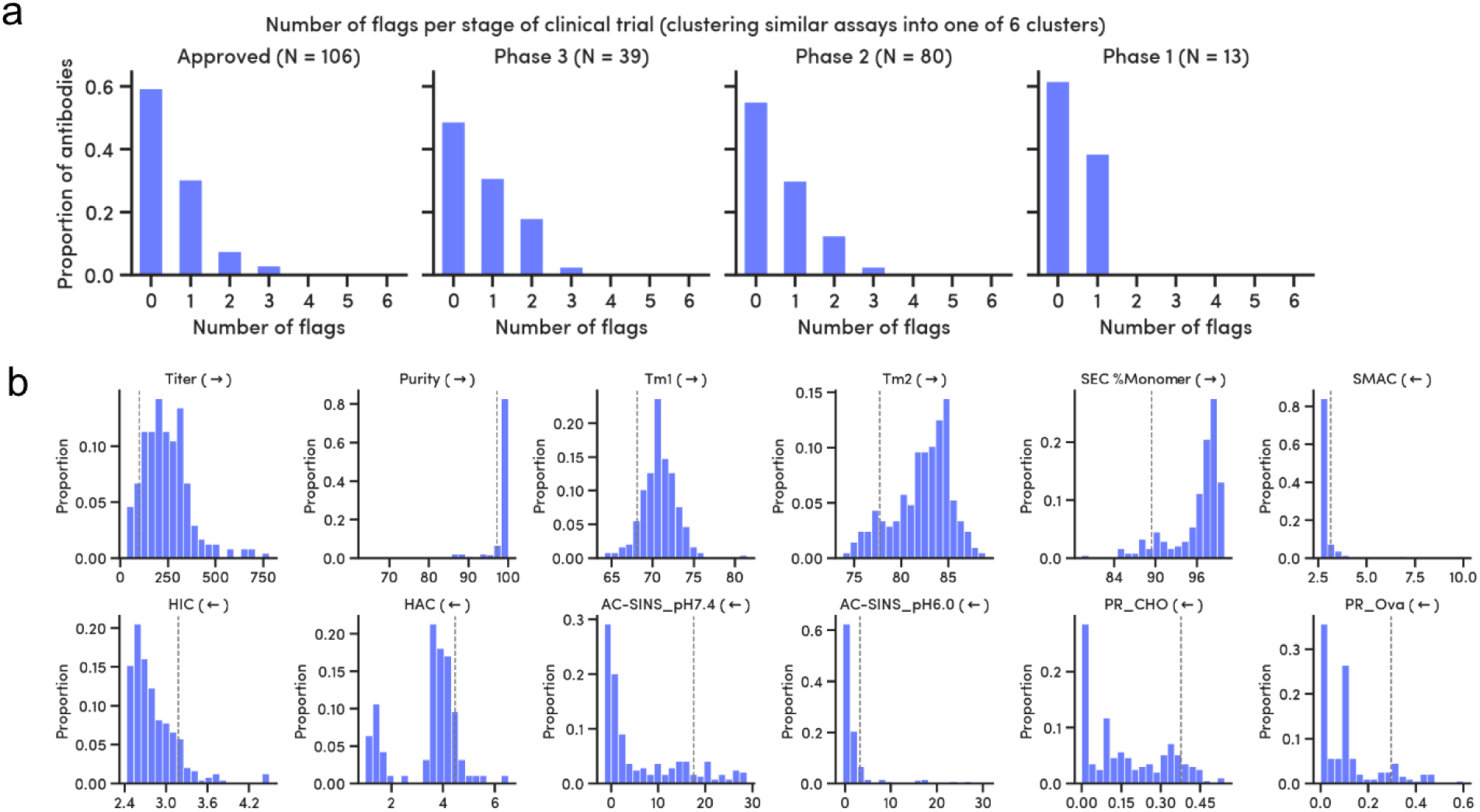
*(a)* Number of threshold flags per antibody, grouped by clinical trial phase. *(b)* Flag cutoff at the 90th percentile of the distribution of each assay result (approved antibodies only). Arrows indicate the favorable direction (e.g., higher purity is better); vertical grey lines denote the “warning threshold.”

**Table 2.** Updated developability warning thresholds. Thresholds were determined using bootstrapping of the 90th percentile value of each assay for the set of approved antibodies. * Equivalent thresholds to previous studies were estimated using a linear fit: SMAC-RT equivalent would be 10.9 ± 0.7 min, HIC equivalent would be 11.4 ± 0.2 min, and HAC (Hep RT) would be 0.81 ± 0.04 min. ** The CHO SMP and Ova assays do not have clear equivalents in previous scales in the Jain papers, but we have shown that they correlate well with the assays proposed in Makowski et al. 2021. Makowski used a different method to calculate thresholds (cut-offs), whereas we recompute thresholds similarly to the previous Jain methodology.

| Assay | Threshold per Jain 2017 | Approved antibodies in 2017 | Threshold per Jain 2023 | Approved antibodies in 2023 | Threshold per Makowski 2021** | Threshold per Ginkgo 2025 | Approved antibodies in 2025 | Units | Threshold directionality |
| --- | --- | --- | --- | --- | --- | --- | --- | --- | --- |
| Tm1 |  |  |  |  |  | 68.09 ± 0.40 | 102 | Temperature (°C) | Lower |
| Tm2 |  |  |  |  |  | 77.71 ± 0.49 | 90 | Temperature (°C) | Lower |
| Fab Tm | 64.9 ± 0.93 | 48 | 64.2 ± 0.81 | 64 |  |  |  | Temperature (°C) | Lower |
| HIC* | 11.63 ± 0.41 | 48 | 11.53 ± 0.27 | 64 |  | 3.18 ± 0.04 | 104 | Retention time (min) | Upper |
| SMAC* | 12.82 ± 1.20 | 48 | 12.62 ± 1.34 | 64 |  | 3.15 ± 0.13 | 104 | Retention time (min) | Upper |
| HAC* | 0.81 ± 0.06 | 44 | 0.84 ± 0.05 | 60 |  | 4.46 ± 0.12 | 31 | Retention time (min) | Upper |
| AC-SINS, pH6.0 |  |  |  |  |  | 3.3 ± 0.77 | 104 | Nanometers | Upper |
| AC-SINS, pH7.4 | 11.78 ± 6.23 | 48 | 13.84 ± 4.80 | 64 |  | 17.51 ± 2.38 | 104 | Nanometers | Upper |
| PSR SMP | 0.26 ± 0.06 | 48 | 0.31 ± 0.05 | 64 |  |  |  | Polyreactivity score (Unitless) | Upper |
| PR CHO** |  |  |  |  | 0.19 | 0.38 ± 0.02 | 83 | Polyreactivity score (Unitless) | Upper |
| PR Ova** |  |  |  |  | 0.11 | 0.30 ± 0.05 | 83 | Polyreactivity score (Unitless) | Upper |

Figure 9a shows the distribution of warning flags for antibodies at different stages of clinical development. We see only a slight decrease in flag count as antibodies advance through clinical development (Figure 9a), an effect much less pronounced than that observed by Jain et al., 2023. We also analyzed the evolution of antibody flag count over the years 2000 to 2015. The flag count does not substantively vary over time, regardless of clinical stage (**Figure S3**).

The flag/threshold-based approach, while offering a simple framework for initial risk assessment, suffers from several limitations. By reducing complex developability data to binary outcomes, it oversimplifies intricate relationships and obscures potentially compensatory effects between different properties. The chosen 90th percentile thresholds are arbitrary, lack contextual information regarding target or indication, and offer limited predictive power for clinical success. Furthermore, the flags and 90th percentile thresholds rely heavily on the representativeness of the training data and ignore the potential for engineering solutions to mitigate identified liabilities. Therefore, while valuable as a starting point, this method necessitates a more nuanced, data-driven strategy to accurately predict developability and clinical outcomes.

### An ML-based Classifier Improves Prediction of Antibody Approval Outcomes

Given the inability of the simple flag-based approach to distinguish clearly between approved and non-approved antibodies (Figure 9), we implemented an ML approach that leverages the full suite of developability data to enable more reliable downselection and facilitate prioritization of candidates with the greatest potential for clinical success.

We trained our ML model to predict antibody approval status (a binary outcome) based on developability assay measurements as the input feature variables. We restricted our analysis to 106 approved and 91 not-approved antibodies, excluding those currently in development. We trained twenty XGBoost classifier models (80:20 train:test split), using bootstrapping to improve statistics given the limited data, and aggregated performance across the 20 models to provide the final classifier. We compared results to 20 null hypothesis models trained on shuffled labels. The results are shown in Figure 10.

**Fig. 10.**
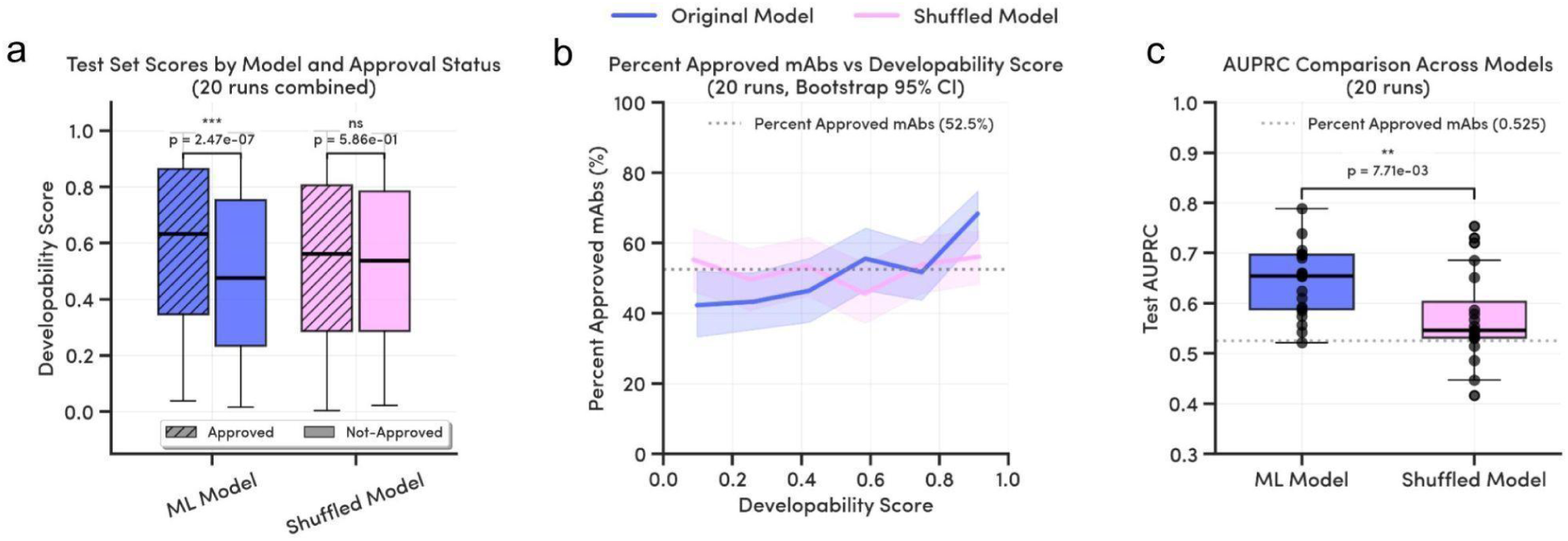
*(a)* Difference between predicted developability scores from twenty bootstrap-sampled XGBoost models compared to twenty models with shuffled labels (null hypothesis); p-value based on the Mann-Whitney U test between “Approved” and “Not-Approved” clinical antibodies. *(b)* Percentage of Approved antibodies as a function of developability score; error bars correspond to 95% confidence intervals from a Wilson score interval for binomial distributions. The dashed line corresponds to the percentage of approved antibodies in the original dataset. (c) Area under the precision–recall curves (AUPRC) for twenty XGBoost models and twenty shuffled label models. The dotted line as in (b).

The ML-model output is the predicted likelihood of approval of a given antibody, which we call the “developability score.” As shown in **Figure 10a**, average developability scores of approved antibodies (0.60) are significantly higher (p-value 3.5e-5) than not-approved antibodies (0.49); as expected, the shuffled (null) model shows no such discrimination. This result indicates that the developability assay data of approved antibodies contain latent features that can indeed be used to predict approval likelihood, even in light of other factors that may have led to antibody discontinuation.

Higher developability scores are more likely to be associated with approved antibodies, as shown in **Figure 10b**. Indeed, the lowest-scoring antibodies are up to 20% more likely to be not-approved, relative to the null hypothesis, and the highest-scoring antibodies are up to 15% more likely to be approved. Lastly, in **Figure 10c**, we compare the area under the precision–recall curve (AUPRC) for the 20 ML models and the shuffled (null) models to give a better sense of overall predictive performance. The median AUPRC is 0.63 for the correct labels, whereas it is only 0.56 for the shuffled labels—right at the null hypothesis positive value rate.

Taken together, these results suggest that our ML-based approach is superior to the flag-based approach in determining the approval likelihood of an antibody based on developability assay assessments. As the developability data have predictive power, we believe our ML-based approach will help guide future antibody discovery campaigns, steering them toward greater clinical success. We expect the power of this modeling approach only to improve as we collect more data on approved and not-approved antibodies in the coming years, furthering the need for developability assay measurements.

### Antibody Developability Predictive Models Improve with More Training Data

We trained predictive antibody developability models using protein Language Model (pLM) embeddings combined with linear regression, a common baseline in the field [Hsu et al. 2022, Notin et al. 2023]. Our goal was to showcase the utility of these data in modeling developability properties, explore and publish results for various model architectures, and garner an understanding of how increased dataset size impacts model performance.

We generated sequence embeddings using ESM-2 [Lin et al. 2023] pLMs, with 8 million (M) to 650M model parameters. We used one-hot encodings of the ANARCI-numbered sequences [Dunbar and Deane 2016] as baseline. pLM embeddings outperformed the baseline (**Figure 11a**), but the number of parameters did not meaningfully nor consistently improve performance distributions. Spearman correlations with ESM-2 (8M) are shown in **Figure 11b**, and predicted values in **Figure S4**.

**Fig. 11.**
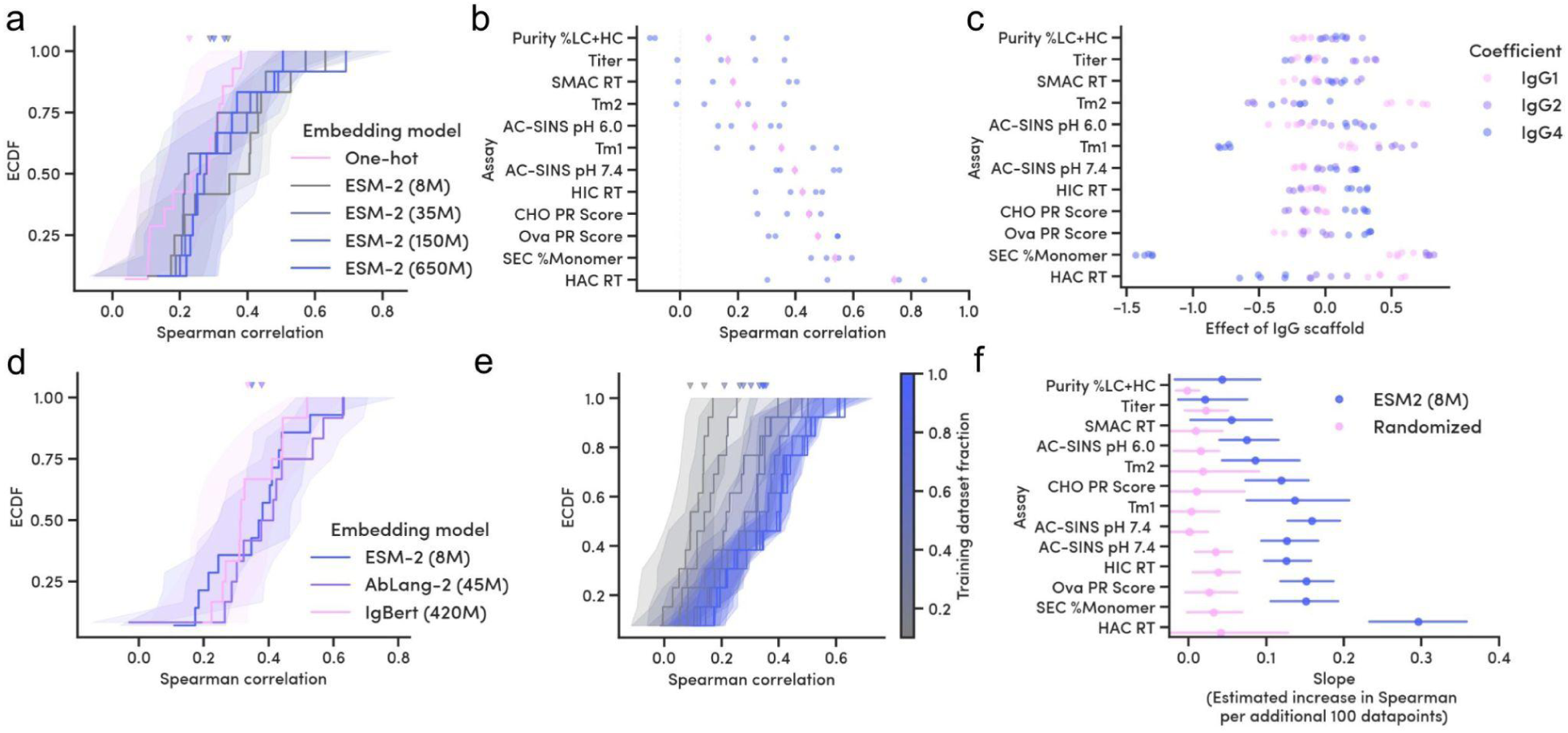
Predictive models based on protein language model embeddings improve with more training data. *(a)* Empirical Cumulative Distribution Functions (ECDFs) of Spearman correlation coefficients between model predictions and measured properties, using embeddings from ESM-2 models or a one-hot encoding. Bootstrapped 95% confidence intervals (CIs; shading); median cross-validation (CV; line); distribution median (triangles). *(b)* Prediction correlations by assay (ESM-2 (8M) embeddings), sorted by median (pink diamonds). *(c)* Linear regression coefficients by IgG subclass (ESM-2 (8M) embeddings or one-hot encoding). *(d)* As in (a), comparing ESM-2 (8M) with antibody-specific pLM embeddings. *(e)* As in (a), comparing ESM-2 (8M) models trained with increasing amounts of data (low (grey) to high (blue)). *(f)* Linear model coefficients fit to Spearman correlations as shown in (Fig. S6), for each assay, with ESM-2 (8M, blue) and shuffled datasets as baseline (pink). Slopes roughly express how much the correlations increase for each additional 100 data points. CIs are from 5 linear models, each fit using a CV fold and varying dataset size.

We included IgG isotype information in our models, represented using one-hot encodings; **Figure 11c** shows the magnitude of the corresponding linear model coefficients. Consistent with our exploratory data analysis above (Figures 3–7), IgG isotype had a meaningful effect on several of the developability property measurements, most notably SEC % monomer and T_m1_.

ESM-2 models are pre-trained on monomeric protein sequences. We tested whether embeddings from two pLMs pre-trained on paired antibodies, AbLang-2 [Olsen et al., 2024] (45M parameters) and IgBert [Kenlay et al., 2024] (420M), improved performance. While embeddings from AbLang-2 marginally outperformed ESM-2, neither of these two pLMs meaningfully improved predictive performance, nor did their larger parameter size (**Figure 11d**).

To assess how the amount of training data influences predictive performance, we simulated smaller training datasets by sampling. The distribution of Spearman correlations consistently increased with dataset size (**Figure 11e**); we found that model complexity, using the number of PCA dimensions as proxy, was not a major confounder in this experiment (**Figure S6**). We quantified this increase by fitting linear models to these correlations (**Figure S5**); for simplicity, we expressed their coefficients as the gain in Spearman correlation per additional 100 datapoints (**Figure 11f**). For most assays, where coefficients were above background, these estimates support that additional data, around the order of magnitude generated in this work, leads to meaningful improvements in predictive performance.

Although we only explored the effects of embedding models and training dataset size on prediction in this proof-of-concept study in a preliminary way, our effort suggests that developability properties, as assayed here, are meaningfully influenced by, and can generally be predicted from, the amino acid sequence of paired antibodies, in particular the variable regions. Moreover, the scale of data collection in this work is sufficient to evince small but quantifiable improvements in predictive performance. We expect that the increased data-generation throughput of the PROPHET-Ab platform will drive the development of improved predictive models, and accelerate data-driven discovery and optimization of antibody therapeutics.

## Conclusions and Future Perspectives

This study presents a comprehensive assessment of antibody developability profiles, across 246 IgGs, using Ginkgo’s HT PROPHET-Ab platform. Our findings highlight the complex interplay of biophysical properties key to successful antibody therapeutics. Our dataset is one of the largest developability datasets, with the most biophysical parameters, now publicly available.

The good run-to-run consistency we observe across many assays underscores the robustness and reliability of the PROPHET-Ab platform (Figure 12). Additionally, our HT methods correlate well with previously established methods, validating the accuracy and quality of our platform.

**Fig. 12.**
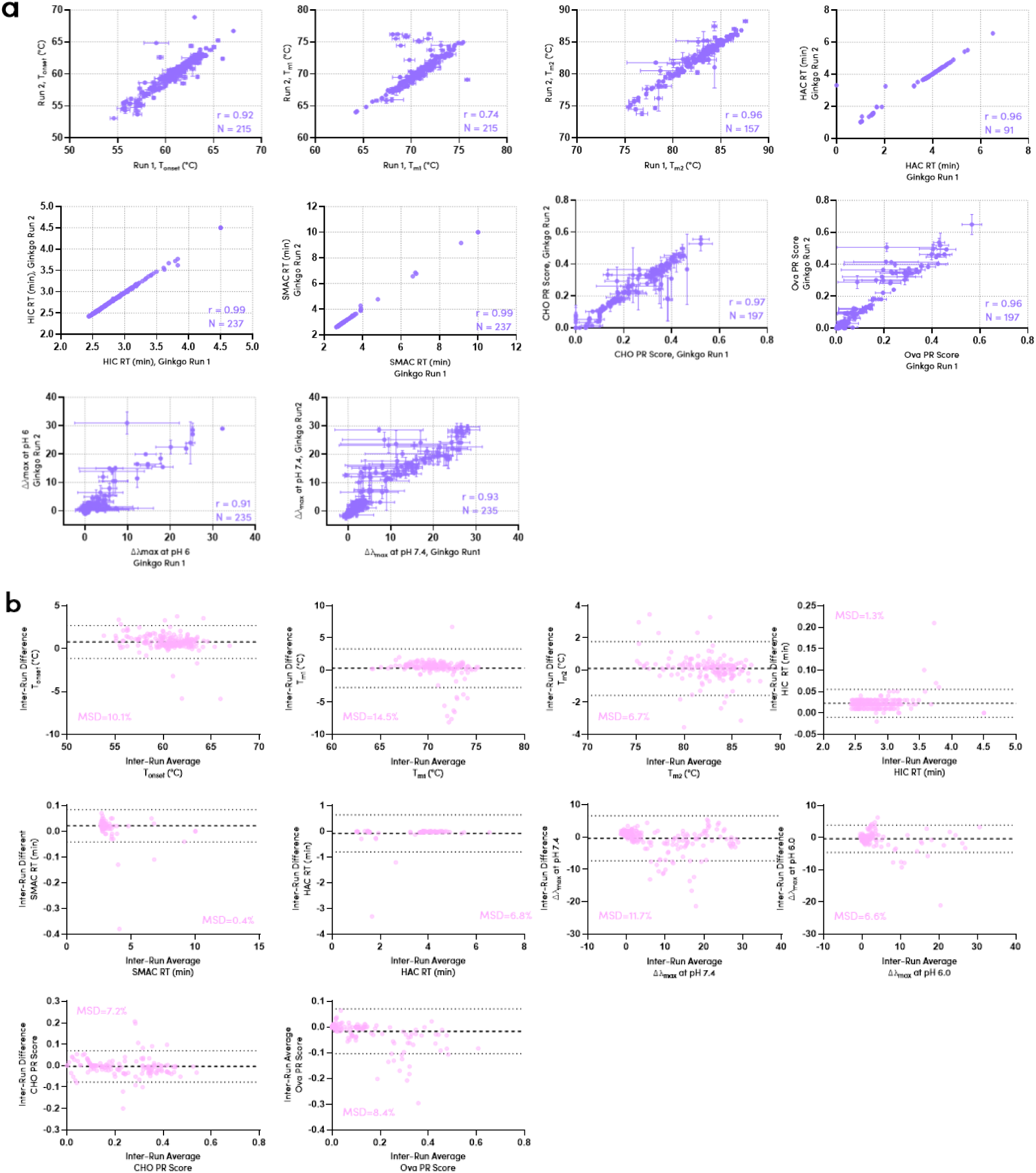
PROPHET-Ab platform robustness, reliability, and quality is demonstrated by high run-to-run consistency across various developability properties (2 independent runs for each assay, each with up to 3 replicates). Data are displayed as *(a)* correlation plots or *(b)* Bland–Altman residual plots.

As the field advances, we anticipate that ever-larger datasets will be needed to fully realize the potential of AI/ML in the prediction of antibody developability characteristics. PROPHET-Ab enables rapid generation of the large volumes of data needed to advance the field, and has the potential to address the current data bottleneck. By automating and optimizing known assays, and integrating them into a robust data analytics pipeline, we have achieved a significant increase in throughput, enabling data acquisition for thousands of antibodies per week. Such an increased volume of data—in a readily ingested “tidy data” format—is crucial for training robust AI/ML models and addresses a key need in biopharma R&D. Indeed, our preliminary tests suggest that model predictive power increases as more training data is used.

PROPHET-Ab enables a virtuous cycle: the generation of more experimental data fuels the development of better predictive models, which are in turn applied to new molecules and new models, accelerating data-and AI/ML-driven antibody discovery and optimization. As the field moves increasingly towards more complex antibody formats such as multi-specifics, we believe the need for HT developability assessment will grow. The PROPHET-Ab platform is well-positioned to meet this demand.

## Materials and Methods

### Antibody Production

The amino acid sequences for the 246 IgGs were extracted from published work (Jain et al., 2017, Ahmed et al., 2021). Antibodies were expressed as their native subclass (IgG1, IgG2, or IgG4), or as IgG1, if a non-standard IgG format. Amino acid and codon-optimized DNA sequences are provided in the Supplementary Information. Antibodies were expressed in HEK Expi293F cells in four separate production batches. Batches 1, 2, 3, and 4 comprise 246, 80, 5 and 5 IgGs, respectively. Batch 3 and 4 antibodies were used exclusively for the DLS-kD assay. Batch 5 comprises 20 IgGs (a subset of the 246 IgGs) that were produced in ExpiCHO cells (sequence data not included). For all batches, standard vendor-recommended expression conditions were used.

*Batches 1 and 3:* VH-encoding sequences were subcloned into the pTwist CMV hIgG1, 2, or 4 expression vectors, and VL-encoding sequences were cloned into pTwist CMV hIgGK or L expression vectors (Twist Bioscience, CA, USA). Heavy and light chain plasmids were transfected in a 1:2::HC::LC ratio in Expi293F cells (Thermo Scientific) as 8 x 1-mL wells of a 96-deep-well plate. *Batches 2 and 4:* VH-encoding sequences were subcloned into the pS-GWZ293-Human IgG1, 2 or 4 expression vectors, and VL-encoding sequences were cloned into pS-GWZ293-Human IgG-CL-Lambda or Kappa expression vectors (Genewiz, Azenta Life Sciences, MA, USA). Heavy and light chain plasmids were transfected in a 1:2::HC::LC ratio in Expi293F cells (10-mL, T125 flasks). *Batch 5:* VH-encoding sequences were subcloned into the pS-GWZCHO-Human IgG1, 2 or 4 expression vectors, and VL-encoding sequences were cloned into pS-GWZCHO-Human IgG-CL-Lambda or Kappa expression vectors (Genewiz, Azenta Life Sciences, MA, USA). Heavy and light chain plasmids were transfected in a 1:1::HC::LC ratio in ExpiCHO cells (Thermo Scientific; 20-mL, T125 flasks).

Antibodies were purified using a single-step Protein A purification, eluted using 50 mM sodium citrate, pH 3.0, and neutralized with 10% v/v 1 M HEPES, pH 8.0. For batches 1, 3, and 5, all antibodies were adjusted to 0.25 mg/mL concentration in 43 mM sodium citrate, 148 mM HEPES, pH 6.0 prior to developability assessments. For batches 2 and 4, antibodies were concentrated and buffer exchanged to the corresponding DLS-kD buffer prior to testing.

### Titer

IgG titers were measured using Valita Titer kits (VAL013, Beckman Coulter Life Sciences) using the vendor-recommended protocol. Clarified supernatants were diluted in fresh cell culture media and added to the assay plate. After 5 min incubation, fluorescence polarization was measured (PHERAstar FSX plate reader, BMG LabTech). Titers were calculated from standard curves of reference IgG (matching heavy chain and light chain types). Titer values are reported as µg/mL. Normalized titers were calculated as final purified yield divided by expression volume, and reported as µg/mL.

### Purity %LC+HC (rCE-SDS)

Purity was determined by reducing SDS capillary electrophoresis (LabChip GXII Touch, high-sensitivity assay protocol). 5 µL of 0.25 mg/ml antibody sample was diluted 1:2 in sodium dodecyl sulfate/dithiothreitol (760518; Revvity) and heated at 95 °C for 10 min prior to separation. Purity (%LC+HC) was measured from the generated electropherograms using the system software.

### Thermostability (nanoDSF and DSF)

nanoDSF was measured on SUPR-DSF (Protein Stable) using the manufacturer’s protocol. 10 µL of 0.25 mg/mL sample was dispensed in a 384 well-plate (HSP3886, Bio-Rad) and centrifuged for 1 min (1000 rpm) to remove air bubbles. Samples were subjected to a 25–95 °C temperature ramp (1.5 °C/min). Thermograms were fitted using the SUPR-Suite software; transition peaks are reported as T_m_. DSF was measured on a CFX Opus thermal cycler (BioRad). 9 µL of 0.25 mg/mL sample was dispensed in a 384 well-plate (951020702, Eppendorf) to which 1 µL of SYPRO Orange dye (S6650, Invitrogen) diluted 1:500 in PBS was mixed.

Samples were subjected to a 25–95 °C temperature ramp (2 °C/min). T_m_ peaks were assigned as the first derivative of the raw data using the Bio-Rad analysis software. A subset of 174 antibodies was used to perform a direct comparison between nanoDSF and DSF at a ramp rate of 2 °C/min (see “nanodsf vs. dsf” sheet in the supplemental data file).

### Size-exclusion chromatography

10 µg sample at 0.25 mg/mL antibody was separated by SEC (XBridge Premier Protein SEC, 186009959, Waters; flow rate, 0.36 mL/min; mobile phase, 200 mM sodium phosphate, pH 7.0). Protein peaks were detected by UV (280 nm) and analyzed using the Chromeleon analysis software.

### Standup monolayer affinity chromatography

The SMAC method reported by Kohli et al., 2015 was optimized for higher throughput. A 2.5 µg sample at 0.25 mg/mL was injected on a Zenix SEC-300 column (213300-4615, Sepax Technologies; flow rate, 0.5 mL/min; mobile phase, 150 mM sodium phosphate, pH 7.0; run time,10 min, corresponding to 2 column volumes). Protein peaks were detected by UV (280 nm) and analyzed using the Chromeleon analysis software.

### Hydrophobic Interaction Chromatography

The HIC method reported by Jain et al., 2017 was optimized for higher throughput. A 2.5 µg sample at 0.25 mg/mL was injected on a Proteomix HIC butyl-NP5 column (431NP2-4603, Sepax Technologies; linear gradient of mobile phase A (100% of 1.8 M ammonium sulfate, 0.1 M sodium phosphate, pH 6.5) and mobile phase B (100% of 0.1 M sodium phosphate, pH 6.5) over 4.5 min; flow rate, 1.5 mL/min). Protein peaks were detected by UV (280 nm) and analyzed using the Chromeleon analysis software. For very hydrophobic antibodies that do not elute within 4.5 min, we assign a retention time of 4.5 min, as did Jain et al., 2017 (25 min in their system).

### Heparin Affinity Chromatography

The HAC method reported by Kraft et al., 2020 was optimized for higher throughput. A 2.5 µg sample at 0.25 mg/mL was injected on a POROS™ Heparin 50 μm Column (4333413, Thermo Fisher; linear gradient of mobile phase A (50 mM Tris, pH 7.4) and mobile phase B (20 mM Tris, 1M NaCl, pH 7.4) over 10 min; flow rate, 1 mL/min). Protein peaks were detected by UV (280 nm) and analyzed using the Chromeleon analysis software.

### Polyreactivity (PR CHO and PR Ova)

We followed the PR CHO SMP and Ova methods as described by Makowski et al., 2021, as modified for use of high-throughput automated workcells. Solubilized membrane proteins (SMP; solubilized with 1% *n*-dodecyl-β-D-maltopyranoside) were prepared from CHO cells as reported. Protein A Dynabeads (10002D; Invitrogen) were coated with antibody (30 µg/ml), then washed with PBS–Bovine Serum Albumin (PBSB). CHO SMP and ovalbumin (A5503; Sigma) polyreactivity reagents were biotinylated (NHS-LC-Biotin; Pierce, 21336; Thermo Fisher). Each biotinylated reagent was incubated with the antibody-coated Dynabeads, washed, and labeled with streptavidin-AF647 (S32357; Invitrogen) and anti-(human Fc) F(ab′)_2_ AF488 (H10120; Invitrogen). Beads were washed again with PBSB and analyzed via flow cytometry (iQue3; Sartorius) to measure median fluorescence intensity (MFI). MFI values were normalized between 0 to 1 based on three reference antibodies exhibiting low (Elotuzumab), medium (Dalotuzumab), and high (Ixekizumab) MFI values using the scoring system reported by Shehata et al., 2019.

### Self-Association (AC-SINS)

The AC-SINS assay was performed as described (Sule et al., 2011, Liu et al., 2014) using automated workcells for higher throughput. Gold nanoparticles (15705; Ted Pella Inc.) were coated with an 80:20 mixture of anti-(human Fc) goat IgG and polyclonal goat nonspecific antibody (109-005-098 and 005-000-003; Jackson ImmunoResearch). Test antibodies were incubated with the nanoparticles in either PBS, pH 7.4 or 20 mM histidine, 150 mM arginine•HCl, 0.05% polysorbate 80, pH 6.0 for 2 h. The wavelength shift was determined from absorbance spectra (PHERAstar FSX plate reader (BMG LabTech), 450–650 nm in 1 nm increments). Samples with a higher degree of self-association show a larger wavelength shift.

### Self-Association (DLS-kD)

DLS-kD was performed on 10 antibodies in PBS, pH 7.4, and on 5 antibodies in 20 mM histidine, 150 mM arginine•HCl, pH 6.0. Antibodies, concentrated to 7.5 mg/ml, were used to generate a six-point concentration curve (7.5, 5, 4, 3, 2, 1 mg/ml) in a 384-well plate (P8806-38403; Waters), centrifuged at 1000*g* for 2 min to remove air bubbles. Samples were analyzed using dynamic light scattering on a Dyna Pro III reader (Waters; 10 acquisitions, 2 sec). The system software was used to calculate kD by plotting the diffusion coefficient against the concentration.

### Data Consolidation and Analysis Data File

The supplemental data file contains:

1. “Definitions” sheet describing each column header corresponding to a data output;
2. “Sequences” sheet documenting the amino acid and codon optimized DNA sequences for all 246 IgGs across the multiple batches produced in this work, the highest clinical status reached by the antibody, as of Feb 2025 as per Thera-SAbDab database, and the current approval status;
3. “Assay Data - tidy format” sheet documents all data generated in this work in a tidy data format (one row per replicate) across 246 IgGs, 4 production batches, up to 3 technical replicates per batch, for 10 developability assays (with 2 conditions each for AC-SINS, DLS-kD, Polyreactivity);
4. “Assay Data - average” sheet summarizing average, standard deviations and number of replicates for each developability assessment (one row per antibody);
5. “nanodsf vs dsf” sheet documenting thermostability data generated with a ramp rate of 2 °C/min (as opposed to the thermostability data in the main sheets, which was generated at 1.5 °C/min) in a tidy data format for 174 antibodies; and
6. “Prior literature Data” sheet summarizing prior published developability assessment data (to perform correlation analysis with data generated in the current work).

Unless otherwise specified, all figures and tables use the data for an antibody averaged across all replicates from both production batches 1 and 2.

Infliximab and Pembrolizumab were only produced in Batch 3 and assayed exclusively for DLS-kD and none of the other assays. **Table S1** provides an explanation for missing datapoints, short of 246, for any assay where applicable.

### Statistical Analysis

Statistical analysis was performed with R (version 4.2.3), Spotfire (version 14.0.5), and Prism (version 10.4.1). Due to the non-normal distributions of some antibody characteristics, Spearman’s rank correlations were calculated for all pairwise combinations. P-values of each correlation were calculated to determine statistical significance (**Table S2**). Inter-run reproducibility was analyzed to determine the linear Pearson correlation, as well as Bland–Altman analysis and mean significant difference (MSD) to determine the agreement.

### Rank Correlation Clustering

We generated a pairwise Spearman rank correlation coefficient matrix for all properties. The pandas DataFrame.corr() function, method=”spearman”, was used to calculate the matrix, which uses pairwise not-a-number (NaN) dropping (generating identical results to the R *cov* method with use=“pairwise.complete.obs”). For hierarchical clustering, the sklearn.cluster.AgglomerativeClustering function was used, with distance=”precomputed”, linkage=”average”, and passing as input 1 – *corr_matrix_abs*, where *corr_matrix_abs* is the absolute values of the pairwise correlation matrix. This method is equivalent to previous methods that used the R *hclust* method with average linkage.

### Thresholds

Thresholds were defined, for each property, as the 90th percentile of a property across a reference set of up to 106 antibodies annotated as “Approved” as of February 2025. Thresholds were estimated using 2,000 bootstrap samples of the reference set. We grouped well-correlated properties to avoid double-counting. These groups were assigned using the rank correlation clustering methods above and are shown in Table S3. A *cluster* violation was assigned if any of the properties in a cluster exceeded the 90% threshold. Only assays with >90 data points were included in the flag analysis; thermostability and titer were excluded, as in prior work.

### Classifier Model Training

Data analysis and modeling were carried out using Python 3.11, scikit-learn, xgboost, and standard scientific libraries. All models were trained on the assay measurements averaged by replicates and batches. The following assay data were used to train the classifier(s): HIC RT, SMAC RT, HAC RT, PR Ova, PR CHO, SEC %Monomer, AC-SINS pH 6, AC-SINS pH 7.4, T_m1_, T_m2_, Titer. Data were standardized and fed directly to the model as a feature vector. The model was trained to predict the status of the 106 approved vs 91 not-approved antibodies. A grid search was performed between XGBoost, LightGBM, RandomForest, Logistic Ridge, Support Vector Classifier, KNN Classifier, and AdaBoost classifiers. It was found rather quickly that no matter the choice of hyperparameter for either method, XGBoost performed the best on a hold-out test set of 20%. From there, a focused set of hyperparameters was chosen for XGBoost including the depth, learning rate, number of estimators, subsampling rate, alpha regularization, and lambda regularization. These were tested by training twenty independent XGBoost classifier models with random train-test splits 80:20 without replacement to get proper statistics on a holdout set. To compare to a random test, 20 independent XGBoost classifiers were trained on the same features but with labels shuffled. The optimal XGBoost model has the following hyperparameter information: max_depth=25, learning_rate=0.25, n_estimators=200, subsample=0.8, reg_alpha=0.1, reg_lambda=0.1.

### Predictive Model Training

Data analysis, manipulation, and modeling were carried out using Python 3.11, PyTorch, and Pytorch Lightning, HuggingFace models, scikit-learn, and standard scientific libraries. pLM embeddings were computed using an Nvidia T4 GPU, with an L4 GPU used for larger-parameter pLMs; all downstream model training and evaluation were performed on CPUs. All models were trained on the data averaged across replicates and batches, for each assay, shared in the “In-vitro measurements - average” sheet. Sequence embeddings were generated for each antibody chain, for all pLM layers. Embeddings from each model and chain along the sequence and layer dimensions were mean-pooled; with single-chain pLMs (ESM-2), per-chain embeddings were further concatenated. pLM embeddings were normalized by removing the mean and scaling to unit variance. Embeddings were reduced to 50 dimensions using Principal Component Analysis with a radial basis function (RBF) kernel, and normalized again. One-hot encodings were generated using ANARCI-aligned heavy and light chain sequences, following the AHo scheme. IgG subclasses, one of IgG1, IgG2, or IgG4, were also one-hot encoded, and concatenated with sequence embeddings or one-hot encodings. We used these representations to train L2-regularized (Ridge) linear regression models, with an alpha=1, on assay values as targets, which were also normalized to zero mean and unit variance. All models were trained and evaluated with 5-fold cross-validation (CV). These CV folds were constructed by sequence clustering using mmseqs2 and IgG subclass stratification, generating groups as dissimilar as possible by sequence identity, while attempting to keep IgG subclass representation uniform across groups. To evaluate the effect of dataset size on model performance, we generated 5-fold CV splits, each with one training and one validation subset, and sampled the training subsets. We sampled without replacement 3 times for each dataset size fraction, from 0.1 to 1 at 0.1 intervals. We used these training dataset samples to train models, and evaluated performance on the corresponding full validation fold, which remained constant across sampled fractions.

## Code availability

Code to reproduce the main computational components of the manuscript is available on github: https://github.com/ginkgobioworks/prophet_ab

---

**Fig. S1.**
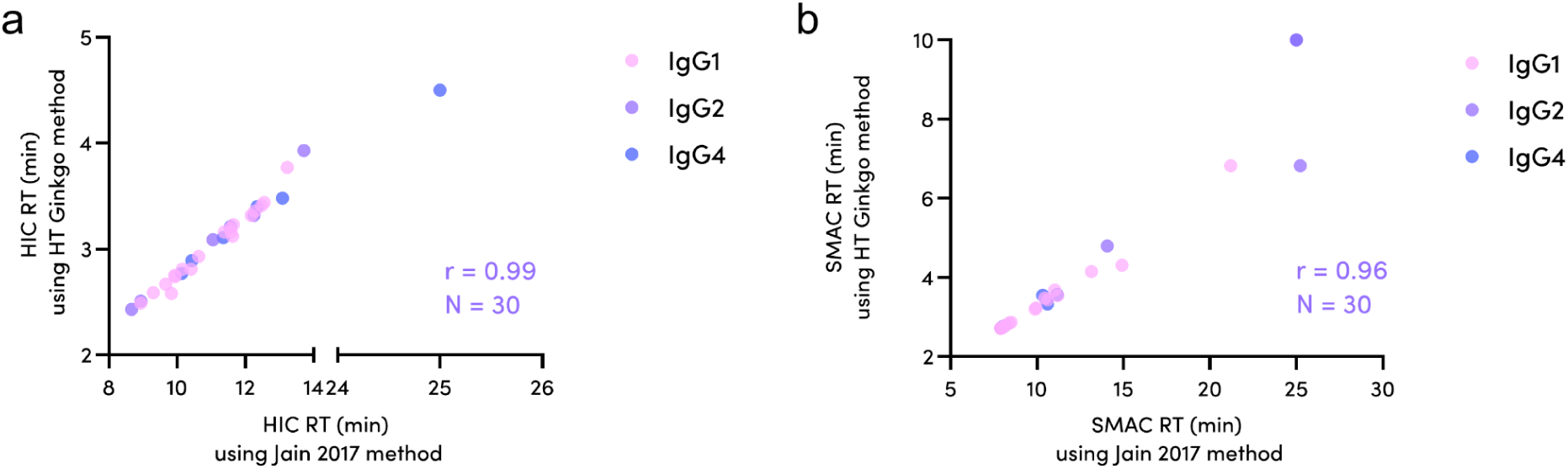
Higher-and lower-throughput *(a)* HIC and *(b)* SMAC data are well-correlated. Ginkgo-generated data, comparing the Ginkgo-developed HT method with the Ginkgo-deployed lower throughput method (Jain et al. 2017) for 30 IgGs chosen for a wide spread in hydrophobicity.

**Fig. S2.**
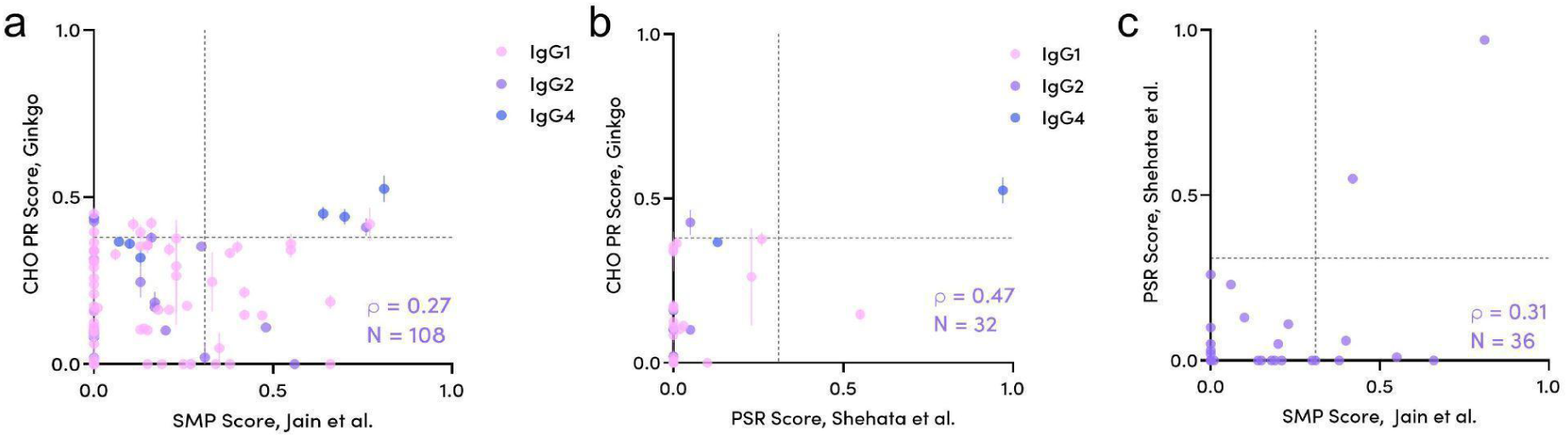
Polyreactivity scores using bead-based methods correlate poorly with yeast-based methods. Ginkgo CHO SMP score compared to *(a)* Jain et al. 2017 SMP (ρ=0.27) and *(b)* Shehata et al. 2019 PSR (ρ=0.47) scores. *(c)* Jain compared to Shehata (ρ=0.31).

**Fig. S3.**
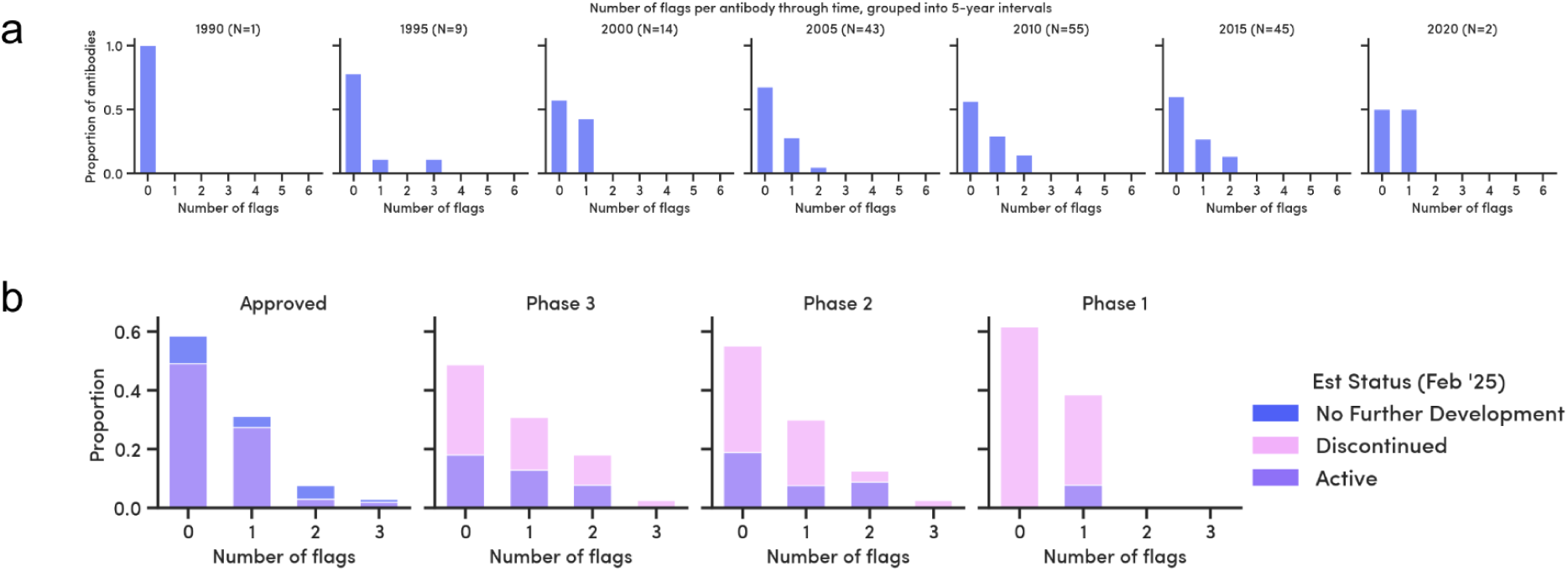
*(a)* Evolution of clinical-stage antibody flag count over time, grouped by the year the therapeutic was proposed. *(b)* Approved or clinical-stage antibody flag counts as of 2025, colored by antibody development status.

**Fig. S4.**
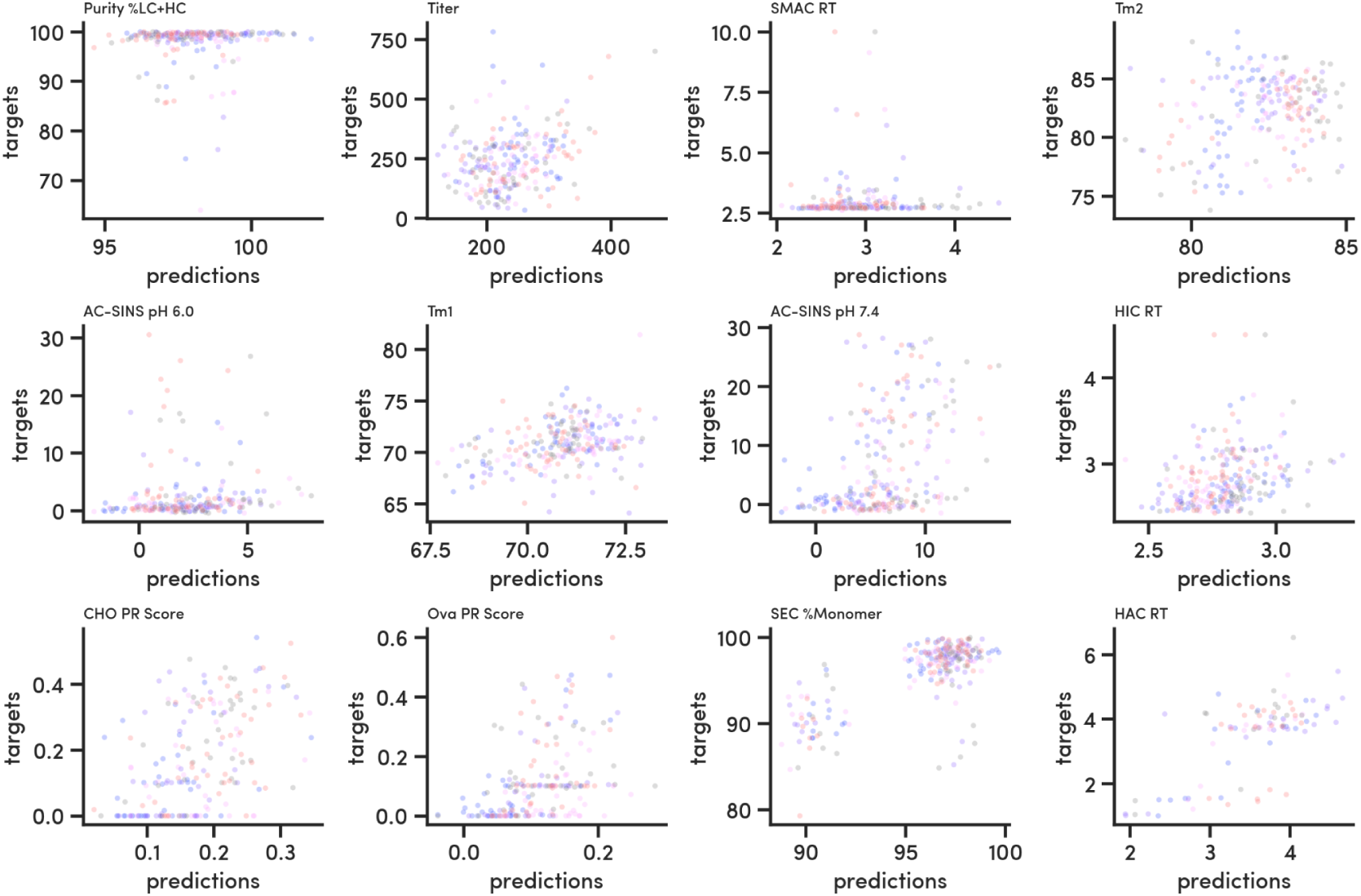
Predicted and measured value pairs from cross-validation by assay, colored by CV fold, using ESM-2 (8M) embeddings and one-hot encoded IgG subclass.

**Fig. S5.**
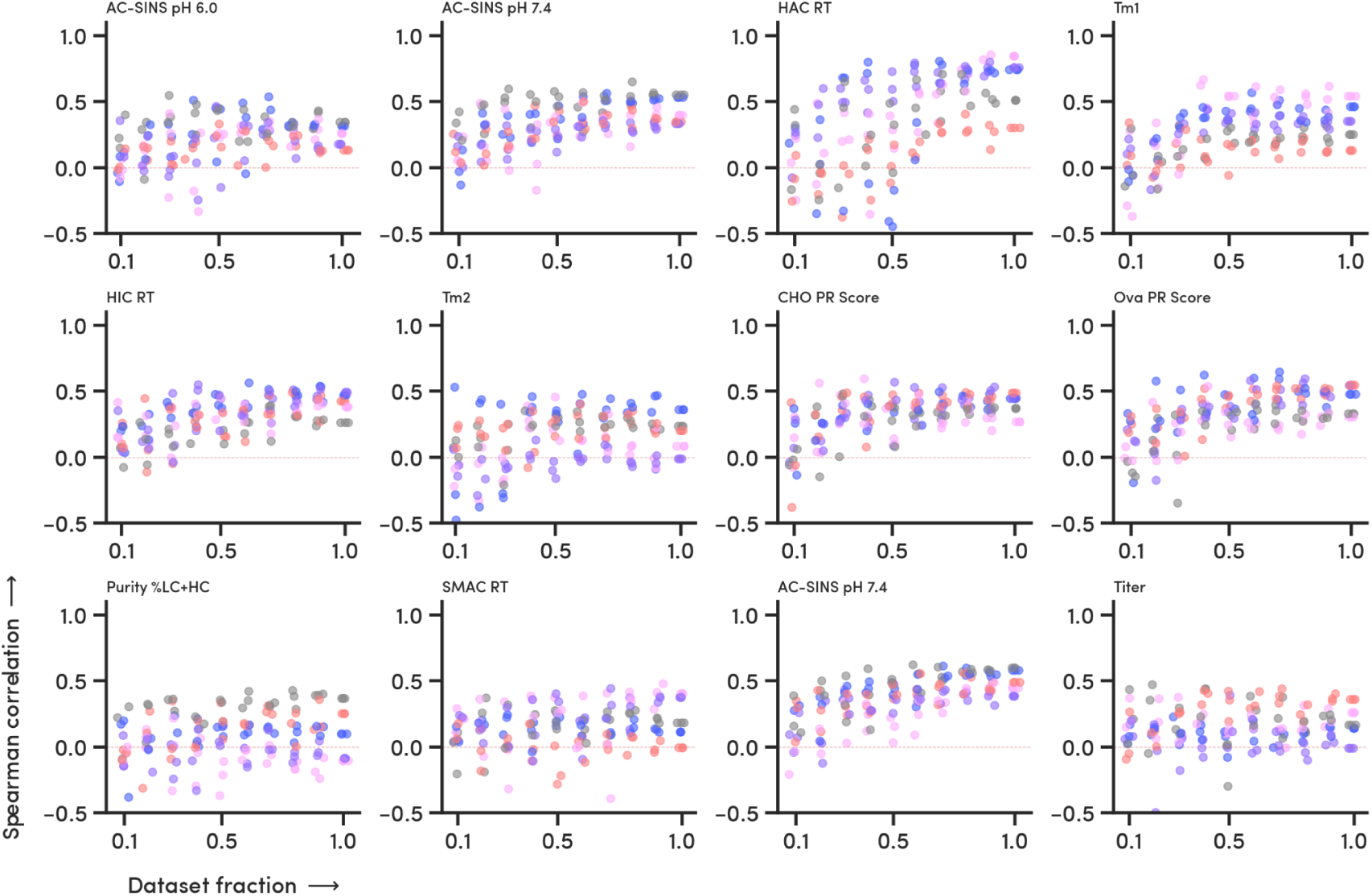
Spearman correlation coefficients for each assay as a function of fraction of the training dataset sampled, colored by CV fold. Each point summarizes predictions from a model trained on a different sample without replacement, using ESM-2 (8M) embeddings.

**Fig. S6.**
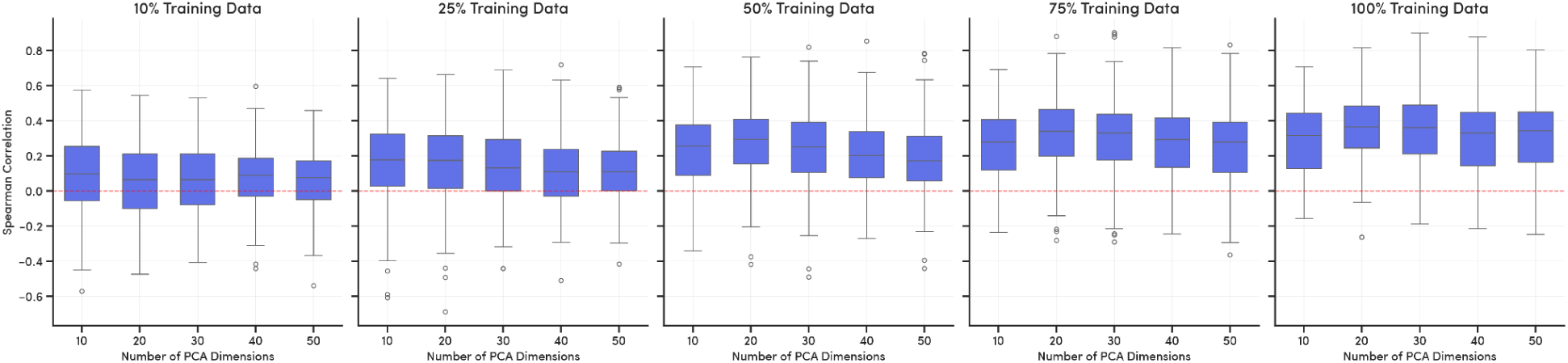
Choice of PCA dimensions in the proposed dataset size and feature size regime does not meaningfully impact the observation of additional training data improving model performance.

**Table S1.** Explanation for missing datapoints for various assays.

| Developability Assay | Related column header/(s) in supplemental data file | Number of missing datapoints | Reason for missing datapoints |
| --- | --- | --- | --- |
| Titer | titer_productionbatch1 | 2 | Data confidence was low due to high variance |
|  | titer_productionbatch2 | 164 | Batch 2, a subset of 246, included only 80 antibodies to study production batch effects |
| Thermostability ($T_{onset}$, $T_{m1}$, $T_{m2}$, $T_{m3}$) | tonset/tm1_nanodsf | 7 | Thermogram fit confidence was low for 7 samples, rendering their data unusable. |
| | tm2_nanodsf | 36 | 29 samples showed a single peak (reported as $T_{m1}$), in addition to the 7 missing datapoints. |
| | tm3_nanodsf | 240 | Most samples showed up to 2 peaks ($T_{m1}$ or $T_{m1}$ & $T_{m2}$) |
|  | tonset/tm1/tm2/tm3_nanodsf_2degC/min_ramp, tonset/tm1/tm2/tm3_dsf_2degC/min_ramp, | 72 | 72 samples were not assayed due to limited yield |
| Polyreactivity (CHO, Ova) | polyreactivity_prscore_cho/ova | 47 | 47 samples were not assayed due to limited yield |
| Heparin binding (HAC) | hac_rt | 150 | 150 samples were not assayed due to limited yield |

**Table S2.** p-values for the correlations presented in Figures 5, 6, and 7.

| Assay 1 | Assay 2 | Number of antibodies | Spearman rank correlation coefficient | p-value |
| --- | --- | --- | --- | --- |
| Ginkgo HIC | Jain et al. 2017 HIC | 133 | 0.97 | <0.001 |
| Ginkgo SMAC | Jain et al. 2017 SMAC | 133 | 0.92 | <0.001 |
| Ginkgo HAC | Kraft et al. 2020 HEP | 49 | 0.92 | <0.001 |
| Ginkgo AC-SINS, pH 7.4 | Jain et al. 2017 DLS-kD | 133 | 0.88 | <0.001 |
| Ginkgo AC-SINS, pH 7.4 | Ginkgo DLS-kD, pH 7.4 | 8 | -0.79 | 0.021 |
| Ginkgo AC-SINS, pH 6 | Ginkgo DLS-kD, pH 6 | 5 | -0.90 | 0.037 |
| Ginkgo PR CHO | Ginkgo PR Ova | 197 | 0.76 | <0.001 |
| Ginkgo PR CHO | Makowski et al. 2021 PSP CHO SMP | 27 | 0.86 | <0.001 |
| Ginkgo PR Ova | Makowski et al. 2021 PSP Ova | 27 | 0.90 | <0.001 |
| Ginkgo PR CHO | Jain et al. 2017 PSR CHO SMP | 108 | 0.27 | <0.01 |

**Table S3.** Group assignments for the cluster flag thresholds.

| Assay | Group assignment |
| --- | --- |
| AC-SINS, pH 6 | 1 |
| AC-SINS, pH 7.4 | 1 |
| PR Ova | 2 |
| PR CHO | 2 |
| HAC | 3 |
| SMAC | 4 |
| HIC | 4 |
| SEC %Monomer | 5 |
| Purity | 6 |

